# Cell atlas of the developing human meninges reveals a dura origin of meningioma

**DOI:** 10.1101/2025.07.08.663122

**Authors:** Elin Vinsland, Sergio Marco Salas, Ivana Kapustová, Lijuan Hu, Simone Webb, Xiaofei Li, Xiaoling He, Mats Nilsson, Muzlifah Haniffa, Roger Barker, Oscar Persson, David R. Raleigh, Erik Sundström, Peter Lönnerberg, Sten Linnarsson

## Abstract

The vertebrate central nervous system is enveloped by the meninges, consisting of the pia, arachnoid, and dura layers. The arachnoid is hypothesised to give rise to the most common primary intracranial tumours, meningiomas. However, molecular evidence supporting this hypothesis is lacking. There are no effective medical therapies to treat meningiomas that are resistant to local interventions, encumbered by our limited understanding of their cellular origin. To address this limitation in our understanding of meningioma biology, we generated a comprehensive reference single cell and spatial transcriptomic atlas of human fetal meninges at post-conceptional weeks 5-13. We found that the meningeal layers develop concurrently, and identified an inner *CDH1*-positive dura cell layer expressing tight junction genes consistent with barrier function. We show that transcriptionally, meningioma cells resemble dura-lineage cells, and that common meningioma driver genes were expressed preferentially in the dura lineage. Our findings suggest that meningiomas originate from dura lineage cells.

**HIGHLIGHTS:**

- scRNA-seq and spatial transcriptomics reveals architecture of human fetal meninges development
- Meningeal layers are formed concurrently, by a gradual refinement of cell states
- A *CDH1*+ inner dura sublayer expresses tight junction genes like the arachnoid barrier
- Meningioma tumours likely originate from inner dura lineage cells, not arachnoid

## INTRODUCTION

The vertebrate central nervous system (CNS) is enveloped by the meninges, comprising the pia, arachnoid and dura matres. The meninges attach the brain to the skull, supply blood flow and immune protection, and provide space through which cerebrospinal fluid flows around the brain. Several CNS developmental defects result from aberrations of the meninges, for example those caused by *FOXC1* mutations^1^. Importantly, the meninges give rise to the most common primary intra-cranial tumours, meningiomas. Although commonly benign or low grade, there are no effective medical therapies for higher-grade atypical or anaplastic meningiomas that are resistant to standard interventions^2^, partly due to our limited understanding of meningioma biology.

Meningioma tumours arise from meningeal cells, but the precise cell type of origin is not fully known. John Cleland (1864) first described meningioma as a tumour of the arachnoid granulations, which are protrusions of the arachnoid mater into the dural venous sinuses that drain cerebrospinal fluid into venous blood. Martin Schmidt (1902) showed that meningioma cells can resemble arachnoid cap cells, which form whorls and psammoma (sand-like) bodies on the perimeter of arachnoid granulations. The influential neurosurgeon Percival Bailey (1931) agreed, noting that meningioma tumours often cause a gritty sound when cut, due to the presence of numerous psammoma bodies. Contemporary authors generally concur with the arachnoid cap cell origin^3^. However, the cell of origin has only been inferred based on anatomical, histological and morphological similarities, while molecular or genetic evidence is lacking. In mouse models, there is evidence that some meningiomas can be generated from *PTGDS*-expressing primordial meningeal cells^4^, but it remains unknown if *PTGDS* identifies a progenitor population in human meninges, or if such cells give rise to meningiomas. Furthermore, recent evidence suggests other cell types could contribute to the earliest stages of meningeal tumorigenesis too, such as *NOTCH3*+ mural cells^5^.

In other tumours, the cell of origin has been inferred from transcriptional similarity to normal cell types, for example based on single-cell atlases^6^. While efforts to map the cellular architecture of the meninges have increased in recent years, few studies have specifically focused on the meninges and meningeal fibroblasts. The first such effort performed single-cell RNA sequencing (scRNA-seq) of embryonic day (E) 14.5 *Col1a1* positive cells in mice, and shed initial light on fibroblast heterogeneity within the pia, arachnoid, and dura maters^7^. This was followed by a scRNA-seq study of the human adult dura^8^, and the adult mouse and human leptomeninges (pia and arachnoid)^9,10^. Other studies incidentally had parts of the meninges in their datasets while sampling the brain^11,12^, or focused their analysis on the cranium^13,14^, or immune cells^15^.

Furthermore, tumours tend to hijack developmental mechanisms^6,16,17^, and the region and severity of meningioma tumours correlate with the embryonic origin of the meninges. The meninges are thought to originate from mesoderm and the neural crest. In accordance, meningioma tumours resulting from *NF2* mutations originate mostly from neural crest-derived convexity meninges, where tumours are also more frequently classified as high-grade compared to mesoderm-derived skull-base meningiomas^17^. Therefore, understanding the molecular determinants of meningeal origin and development are key to identify the drivers of growth and transformation of meningiomas. Despite the clinical importance of meningeal origin for meningioma biology, no comprehensive cell atlas has been published covering the developing human meninges. As a result, not much is known about how the human meninges develops either, such as the genesis of meningeal layers by fibroblasts, their maturation over developmental timepoints, and in co-development and differentiation of other cells in the meninges, like immune cells and vasculature.

Here, we therefore set out to survey the development of human meninges, from embryonic post-conceptional week (PCW) 5-8 and fetal PCW 9-13 (hereafter PCW 5-13 collectively referred to as ‘fetal’), by scRNA-seq and probe-based *in situ* gene expression analysis (Xenium, hereafter ‘spatial transcriptomics’; Fig. 1A). We also collected published bulk RNA-seq and scRNA-seq data for human meningiomas^18^, and generated new spatial transcriptomic data for seven such tumours (Fig. 1B).

**Figure 1.**
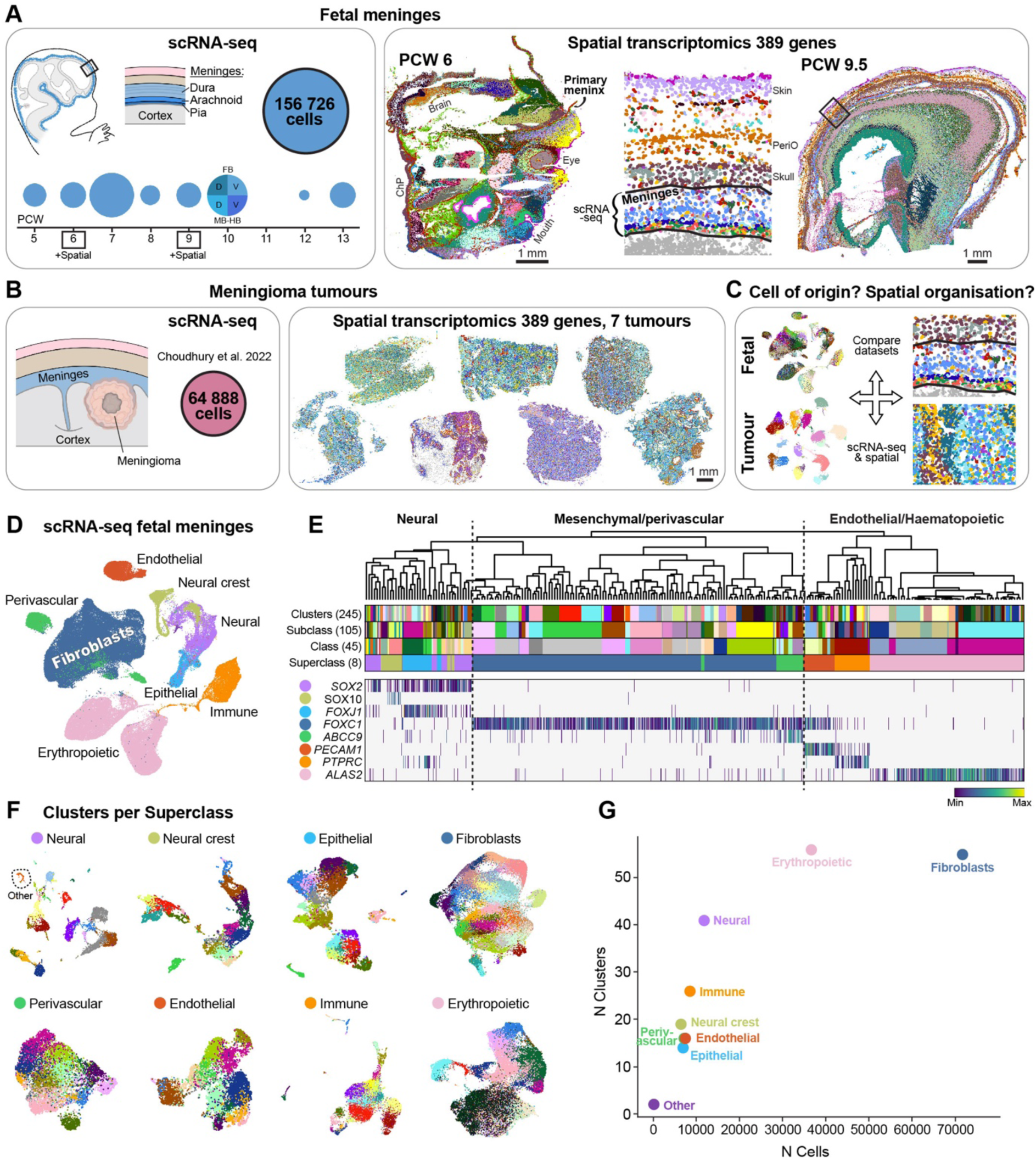
Cellular and spatial atlas of the developing human meninges. (A) Data from fetal meninges. Left: Number of cells collected from PCW 5-13 meninges for scRNA-seq (totally 156,726 cells passing QC). Right: Spatial transcriptomics of sagittal sections of PCW 6 and 9.5 heads. Polygons are segmented nuclei coloured by clusters. Lines define the meninges and where scRNA-seq data was collected. (B) Data from meningioma tumours. Left: scRNA-seq data downloaded from Choudhury et al. (2022)^18^. Right: Spatial transcriptomics performed in this study on seven tumours. (C) Strategy to compare scRNA-seq and spatial data from fetal meninges and meningioma tumours, to study conserved developmental cell types in meningioma. (D) UMAP embedding of scRNA-seq data from fetal meninges coloured by Superclass. (E) Dendrogram of scRNA-seq data with attributes from top to bottom: 245 clusters, 105 Subclasses, 45 Classes, and 8 Superclasses; coloured as in (D), heatmap showing gene expression marking neural cells (*SOX2*), neural crest (*SOX10*), ependymal cells (*FOXJ1*), meningeal fibroblasts (*FOXC1*), pericytes (*ABCC9*), endothelial cells (*PECAM1*), immune cells (*PTPRC*), and erythropoietic cells (*ALAS2*). (F) UMAPs of each Superclass coloured by clusters. (G) Scatter plot of number of cells and number of clusters per Superclass.

Below, we carefully and comprehensively describe our cell atlas of the developing meninges, focusing on immune cells, vascular cells, and importantly, fibroblasts. We then use well-annotated fibroblast cell types and gene expression programs that are active during normal meningeal development as a blueprint to understand the cell of origin and cellular composition of human meningiomas (Fig. 1C).

## RESULTS

### Comprehensive cellular and spatial atlas of the developing human meninges

For scRNA-seq, we collected 16 human meninges samples from 13 donors across PCW 5-13. We collected the whole meninges that peeled off the brain (SFig. 1A-B). A total of 156,726 high-quality cells were used for further analysis (Fig. 1A). The mean UMI per sample was 8,842. As expected, anucleated erythropoietic cells showed the lowest gene and UMI counts (SFig. 1C); excluding those cells, the mean UMI count was 10,249 (Table S1). An analysis of the distribution of maternal and paternal SNP alleles^19^ (Methods) indicated that some erythrocytes were maternal, while all other cells were fetal (SFig. 1D). We found that the data was well-integrated across samples even in the absence of batch correction, which we therefore omitted in order to avoid suppressing true biological variation (SFig. 1E-F).

We generated a total of 245 clusters (Methods), annotated to four levels of detail: cell type (n=115), subclass (n=105), class (n=45), and superclass (n=8). We computed metadata such as cell cycle score, fetal age, and enriched genes for each cluster (Table S2). Since the meninges are a connective tissue, the biggest superclass consisted of *FOXC1*+ fibroblasts, which made up half the dataset. We also identified *PECAM1*+ endothelial, *ABCC9*+ perivascular, *PTPRC*+ immune, and *ALAS2*+ erythropoietic cells in the meninges. Probably, by peeling the meninges off the brain for sequencing, we also acquired interacting *SOX2*+ neural, *SOX10*+ neural crest and *FOXJ1*+ epithelial (choroid plexus) cells (Fig. 1D-G). Due to the diversity of the dataset presented here, we were able to generate highly specific markers for many cell types of the meninges (SFig. 1G).

We used spatial transcriptomics to validate and locate cell types identified in the scRNA-seq data. To keep the meninges anatomically intact we cryosectioned whole heads at PCW 6 and PCW 9.5. Two medio-lateral sagittal sections from PCW 6 and one lateral sagittal section from PCW 9.5 were analysed, with a custom probe-set targeting 389 genes (Table S3) selected based on the fetal scRNA-seq data. We generated 53 and 50 clusters respectively, after single nucleus segmentation at the two ages (SFig. 1H-I, Table S4). At PCW 6, we found diverse populations of craniofacial mesenchyme, neural crest-derived tissues, progenitors and differentiating neural cells of the CNS (in the forebrain, midbrain, hindbrain, eye), choroid plexus, and vascular and immune cells. Importantly, we identified the earliest described layer of the meninges; the primary meninx (SFig. 1H). At PCW 9.5, clustering highlighted a remarkably complete stack of layers spanning the developing cortical layers from the ventricular zone to Cajal-Retzius cells in the cortical plate, to fibroblasts of the meninges, future skull, periosteum, skin and surface periderm (more in Figure 4). We also noted scattered populations of immune cells (for example active and classical monocytes, macrophages, mast cells, microglia), Schwann cells (melanocytic lineage), and various vascular cells, among others (SFig. 1I).

In the following sections, we analysed each superclass of meningeal cells — immune, vascular (Box 1), and fibroblasts — focusing on their heterogeneity and maturation during meninges development.

### Fetal immune heterogeneity and B-lineage cells from PCW 5

The fetal meninges harboured a surprisingly heterogeneous population of immune cells across PCW 5-13, including at the earliest stages. We identified 19 cell types of myeloid and lymphoid origin (Fig. 2A-B, SFig. 2A). Except for three cells, all immune cells were of fetal origin (Fig. 2C). By including canonical markers and enriched genes from our scRNA-seq dataset (Fig. 2E, Table S2) in the spatial transcriptomics panel (Table S3), we validated and located those immune cells in the primary meninx at PCW 6, and leptomeninges and dura at PCW 9.5 (Fig. 2D-G).

**Figure 2.**
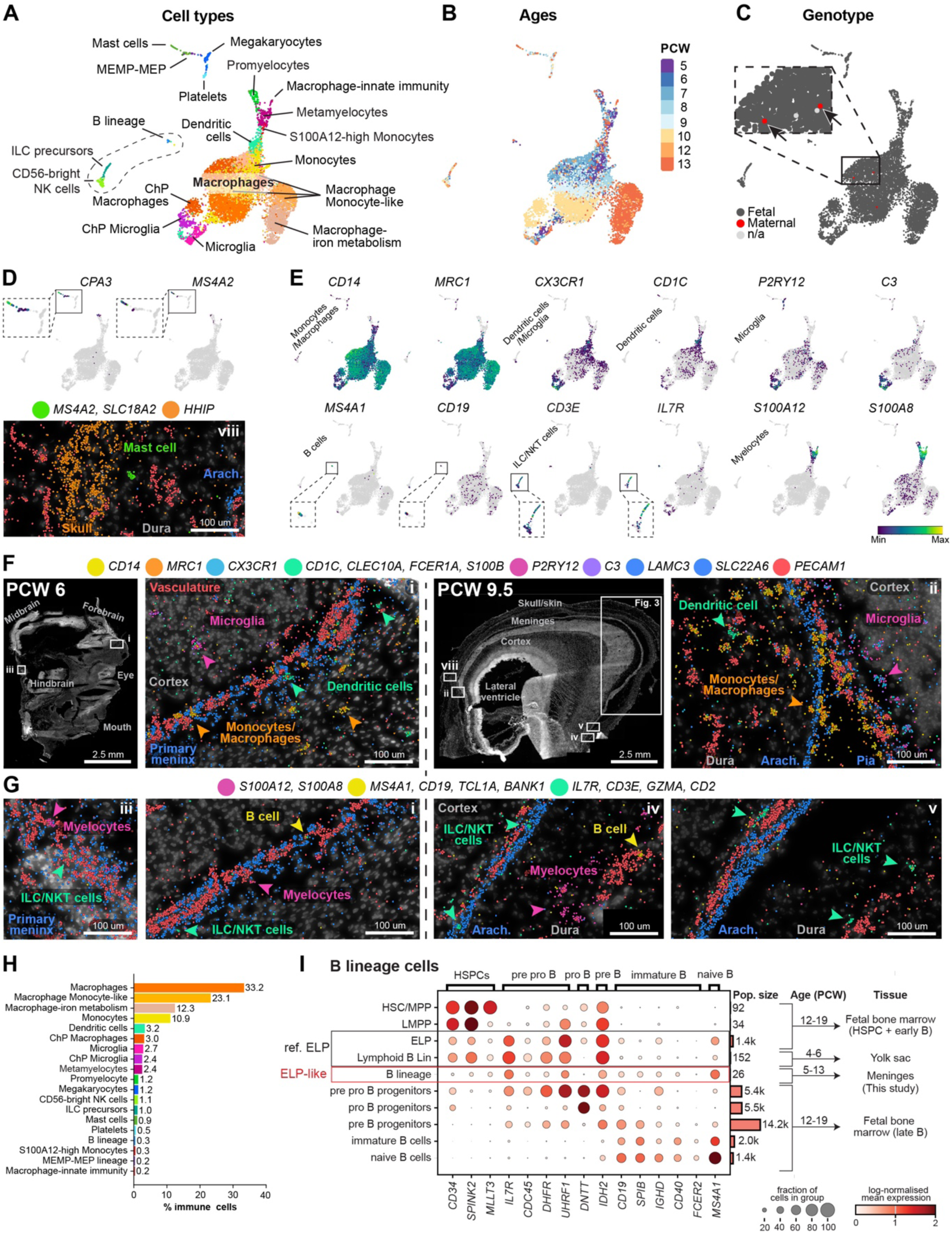
Immune cell diversity in the developing meninges. (A) UMAP coloured by clusters of annotated immune cells. MEMP, megakaryocyte erythroid mast cell progenitor; MEP, megakaryocyte erythroid progenitor. (B) UMAP coloured by sample age. (C) UMAP coloured by maternal/fetal genotype, by maternal/paternal SNP analysis (Methods). Inset highlights two maternal immune cells (arrows). (D) UMAPs coloured by gene expression on a grey background of all cells, and spatial transcriptomics at PCW 9.5, of mast cell markers. (E) UMAPs coloured by gene expression of immune cell markers. (F) Spatial transcriptomics of PCW 6 and 9.5 fetal heads, showing RNA molecules on DAPI. Marker genes for microglia, monocytes/macrophages, and dendritic cells are shown, coloured as in (A). (G) Same as (F) but for ILC/NKT cells, B-cells, and the myelocytic lineage. (H) Percentage of immune cell types, coloured as in (A) (I) Dot plot of scRNA-seq gene expression of B-lineage markers across developmental tissues (fetal bone marrow as B-cell reference^22^ vs yolk sac^21^, and meninges (this data)). The size of the dot represents the % of cells within a group that express a given gene, and the colour of the dot indicates ln(x+1) mean gene expression within the group, where x equal counts normalised to 1e4 per cell. ELP, early lymphoid cell; HSCs, haematopoietic stem cell.

Macrophages were the most abundant (Fig. 2H), with plentiful IBA1+ cells concentrated around the primary meninx/leptomeninges at PCW 6 and 9.5 (SFig. 2B). Notably, IBA1 staining of meninges revealed a uniform sparse tiling of macrophages in en face floating sections (SFig. 2C), consistent with repelling mechanisms controlling their location. The leptomeninges defined an anatomical barrier where microglia (*CX3CR1*, *P2RY12*, *C3*) always resided on the brain side, while monocytes/macrophages (*CD14*, *MRC1*) resided in the leptomeninges and the rest of the head, already from PCW 6 in relation to the primary meninx (Fig. 2F). Microglia were also concentrated inside the developing eye, while monocytes/macrophages resided outside, except one macrophage inside a ring of endothelial cells at the very centre (SFig. 2D).

There is growing evidence for B lymphoid cell presence in early human development. Recent investigations into the fetal B-cell developmental hierarchy defined *CD34*+*IL7R*+*CD19*-early lymphoid progenitors (ELP) in human fetal liver as early as PCW 6^20^. Another report detailed the yolk sac ‘Lymphoid B lineage’ from PCW 4-6^21^. These data raise the intriguing possibility that adaptive lymphoid cells are generated and established in peripheral organs as early as PCW 6. However, the earliest timepoint of establishment of lymphoid cells has not been determined in the embryonic head and meninges. B-lineage cells made up 0.3% of immune cells in the meninges (Fig. 2H) and were clearly distinct from innate lymphoid subtypes (ILC precursors and CD56-bright NK) (SFig. 2E-F). Our scRNA-seq data confirmed that *CD34*+*IL7R*+*CD1*9-low ELP-like B-lineage cells were transcriptionally similar to second trimester fetal bone marrow ELPs^22^, where B cells are well-described to differentiate and diversify (Fig. 2I). We confirmed ELP presence in embryonic meninges as early as PCW 5, and up until about PCW 7, after which point (PCW 8-13) the meningeal B-lineage appeared to differentiate and acquire more mature markers for pre pro-, pro-, pre-, and immature B cells (SFig. 2G).

These data position the meninges as a contributor to the first and second-trimester differentiation and diversification of human B cells, which has recently been appreciated as present in the fetal liver^23^, and a key function of the fetal bone marrow^22^. Further, we demonstrate B cell presence in the embryonic meninges one week earlier in gestation than previously reported for liver tissue.

##### BOX 1 Fetal meningeal vasculature and choroid plexus

###### Meninges contribute to brain vascularisation, and have early PROX1+ lymphatic-like endothelial cells

The fetal meninges contained perivascular cells such as pericytes, smooth muscle cells, and perivascular fibroblasts (Fig. 3A), as well as vascular endothelial cells (Fig. 3B), across PCW 5-13 (Fig. 3C). The endothelial cells expressed stereotypical genes involved in arterio-venous zonation (Fig. 3D-E), as previously described in both mouse and human^24–27^, and also contained *COL15A1*+ *ESM1*+ tip cells (Fig. 3D). Over time, endothelial cells transitioned from the primary vascular plexus at PCW 5, to arterial and venous capillaries around PCW 7-9, and larger arterioles and venous lymphatic-like cells at PCW 12-13. However, the forebrain didn’t have venous capillaries even at PCW 13 (Fig. 3E), indicating that this vascular specification occurred earlier in the meninges. Immunostaining of smooth muscle actin (*ACTA2* gene) showed that larger arteries were present in the leptomeninges, dura and skin at PCW 9.5 (Fig. 3F-G).

**Figure 3.**
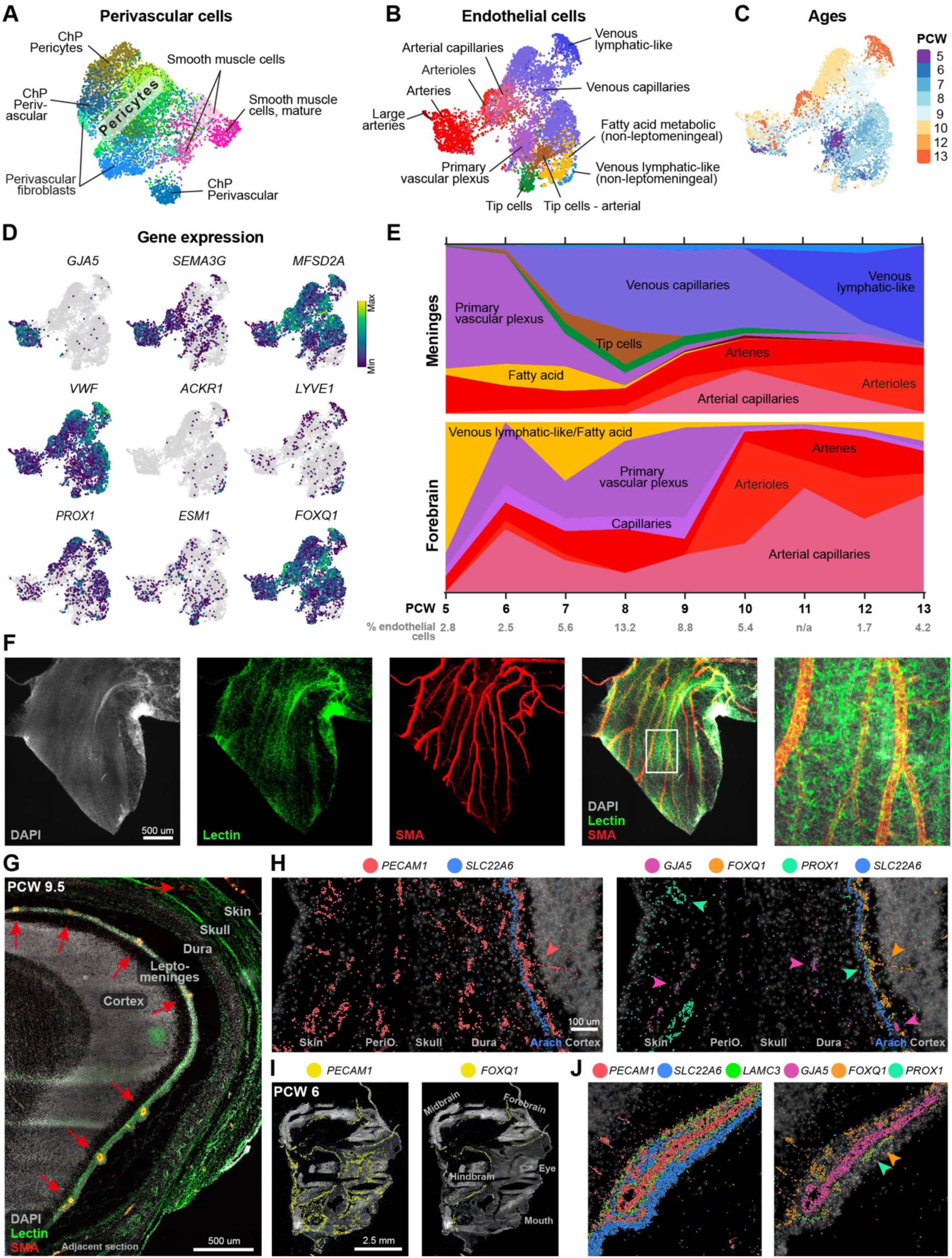
Vasculogenesis in the meninges and brain. (A) UMAP of perivascular cells. (B) UMAP of endothelial cells, coloured by arterio-venous zonation. (C) UMAP of endothelial cells coloured by sample ages. (D) Gene expression of arterio-venous zonation markers, lymphatic- and tip cells. (E) Proportion of endothelial cell types by age, in the meninges and forebrain12. (F) Lectin dye (vasculature), DAPI (nuclei), and Smooth muscle actin (SMA) immunostaining in a PCW 9.5 floating meninges. (G) Same as (F) but in a sagittal section (red arrows point to arteries). (H) Spatial transcriptomics of all endothelial cells (PECAM1), arteries (GJA5 & pink arrows), venous lymphatic-like cells (PROX1 & mint-green arrows), leptomeninges and brain-specific endothelial cells (FOXQ1 & orange arrows) and arachnoid (SLC22A6) in a PCW 9.5 head. Coloured dots represent RNA molecules. (I) Spatial transcriptomics of PECAM1 and FOXQ1 in a PCW 6 head. (J) Same as (H) but with pia (LAMC3), around an artery.

Most endothelial cells in the meningeal scRNA-seq dataset expressed the transcription factor *FOXQ1* (Fig. 3D), which controls differentiation of blood brain barrier (BBB) endothelial cells in mice^28^. Spatial transcriptomics showed that while *PECAM1* was expressed in all endothelial cells in the head, *FOXQ1* was uniquely expressed in endothelial cells below the arachnoid, and extending into the brain (Fig. 3H). This was observed as early as PCW 6 and together with the primary meninx (Fig. 3I). The cellular ontogeny of meningeal lymphatic vessels is still unclear^29^. We found that *ACKR1*+ *LYVE1*+ venous cells expressed the key lymphatic transcription factor *PROX1*^30–32^ (Fig. 3D). Spatially, *PROX1+ FOXQ1*+ endothelial cells were present below the arachnoid, and *PROX1+ FOXQ1*-endothelial cells in the developing skin at PCW 9.5 (Fig. 3H & 3J). These observations suggest that (1) meningeal lymphatic cells were mostly derived from the venous endothelium, and (2) subarachnoid lymphangiogenesis may have started in early fetal development, surprisingly so, since meningeal lymphatics develop postnatally in mice^33^.

###### Choroid plexus ciliogenesis and unique PTPRC+ fibroblasts

CSF is generated by filtration of blood at the choroid plexus (ChP), filling the brain ventricles and subarachnoid spaces. Using spatial transcriptomics, we noticed that three layers comprising the *FOXC1*+ primary meninx, the underlying endothelial cells, and the *FOXJ1+* ependymal cells were continuous with the corresponding three layers of the emerging ChP at PCW 6 (SFig. 3A). At PCW 9.5, the characteristic inside-out anatomy of the ChP had been formed from those initial three layers, with a ChP epithelium surrounding a stroma of meningeal and vascular cells (SFig. 3B-D). There were also erythropoietic cells, immune cells, and perivascular cells in the ChP. The meningeal fibroblasts in the ChP were mostly pial (SFig. 3C). ChP endothelial cells were mostly *FOXQ1*- (i.e. outside brain and leptomeninges), though there were a few *FOXQ1*+ cells (brain-leptomeningeal BBB endothelial cells) too (SFig. 3E). Microglia always resided in the ChP epithelial layer, whilst macrophages were in the stroma, just like the brain versus meninges (SFig. 3F).

The ChP epithelium develops by differentiation from adjacent neuroepithelium (future multiciliated ependymal cells). Our scRNA-seq data comprised 22 clusters of brain and ChP epithelial cells from PCW 5-13 (SFig. 3G-H). While AQP1 immunostaining showed positivity in both brain ependymal cells and the ChP epithelium (SFig. 3D), the serotonin receptor gene *HTR2C* was only expressed in the ChP (SFig. 3I). At the junctions between future ChP epithelium and surrounding neuroepithelium, we identified similar dual neuro-epithelial progenitors and ciliogenesis programs as previously described in mice^34^. These progenitors expressed *RSPO2* and *RSPO3*, followed by *DEUP1* (mouse *Ccdc67*) and *SHISA8* in ciliogenic epithelia, and *AQP1* in fully differentiated ChP epithelia (SFig. 3J-K). We also analysed ChP epithelial cells functionally, by creating a combined CSF gene-set score based on two proteomics datasets from human adult CSF^35,36^ (Table S5). CSF genes were indeed expressed in the fetal data, this being greatest in differentiated ChP epithelial cells (SFig. 3L).

We also identified a fibroblast type only present in the ChP, which uniquely expressed *TCF21* (SFig. 3M-N). Both scRNA-seq and spatial transcriptomics showed that they also expressed the immune-related genes *PTPRC* (CD45, pan-leukocyte marker), and *CD38*. However, mouse *Tcf21*+*Cd38*+ fibroblasts did not express *Ptprc* (SFig. 3M).

Our findings demonstrate that human ChP development uses mechanisms that are largely conserved from rodents, but with human-specific features that suggest an evolved role for ChP fibroblasts in immune surveillance at the blood-CSF border.

### Fibroblast layer development in the fetal meninges and head

Fibroblasts are the principal cells of connective tissues, including the meninges and head mesenchyme. Fibroblasts of the human fetal meninges have previously been described based on basic anatomical, histological and morphological features. However, a comprehensive molecular analysis of head fibroblasts, and the transcriptional cell-type composition of newly forming meningeal layers remain to be elucidated.

### Fibroblast heterogeneity and organisation in the fetal meninges and head

Our scRNA-seq analysis across the PCW 5-13 meninges yielded 71,657 fibroblasts. From this, we generated 55 clusters that we annotated as 29 cell types with distinctly enriched genes (Fig. 4A-B, SFig. 4A, Table S2). 20 of these cell types were classified as fibroblasts of the three meningeal layers; the pia-, arachnoid- and dura mater, their precursors, and the early primary meninx. We also captured other interesting fibroblasts, such as a fibroblast expressing *TAGAP* (a T-cell activating protein), osteogenic fibroblasts (*PTHLH*), chondrocytes (*MATN4*), hindbrain (*HOXA3)* fibroblasts, and the previously mentioned *PTPRC*+ fibroblast unique to the ChP (Fig. 4B-C).

**Figure 4.**
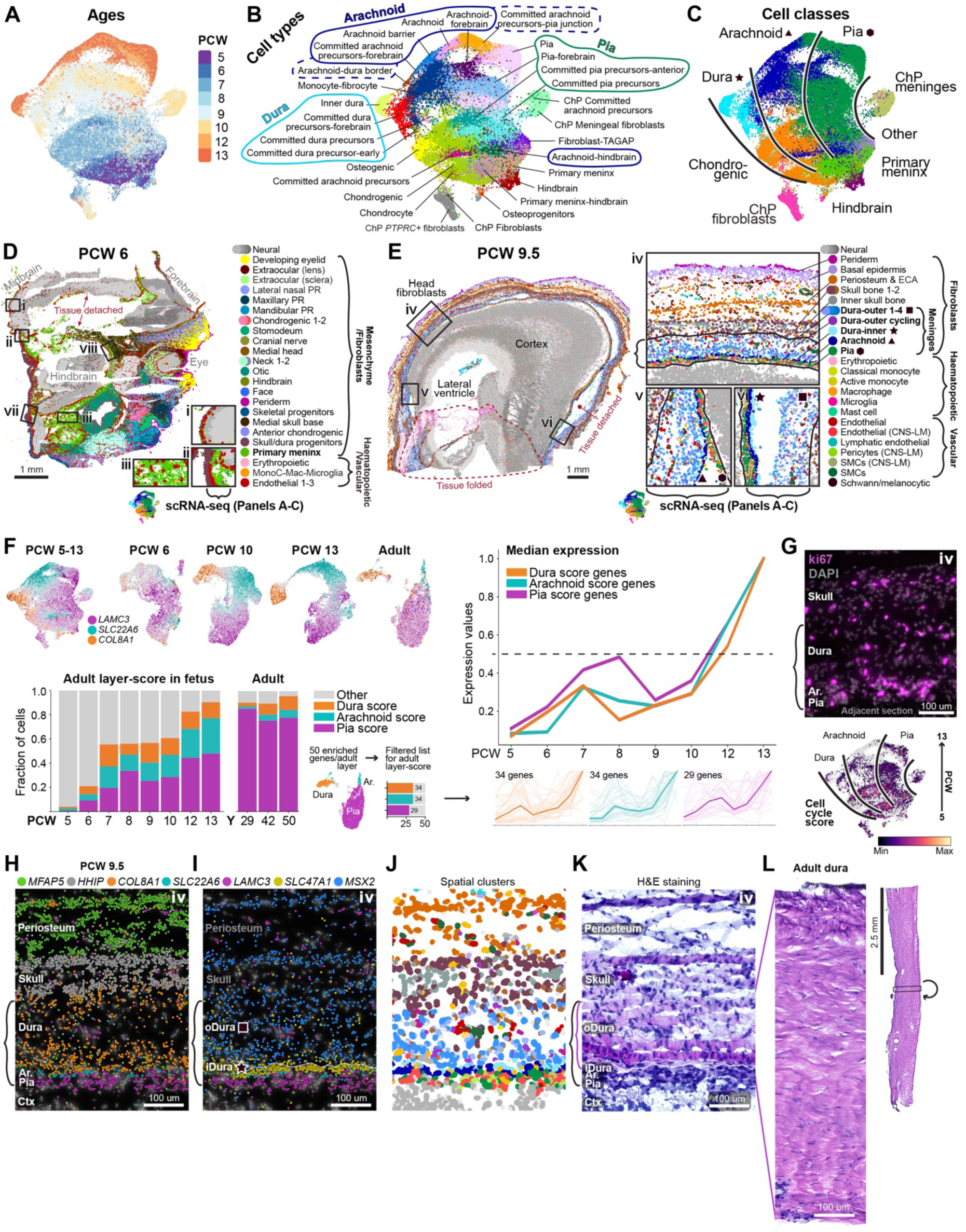
Fibroblast layer development in human fetal heads. (A) UMAP of fetal meningeal fibroblasts coloured by sample age. (B) UMAP coloured by annotated cell types. (C) UMAP coloured by Class. Black lines separate putative pia, arachnoid, and dura lineages. (D) Sagittal section of a PCW 6 head coloured by spatial clusters. Three insets show the primary meninx, and where scRNA-seq was sampled. The colour for the primary meninx matches the scRNA-seq Class ‘Primary meninx’ in (C). Mac, macrophage; MonoC, monocyte; PR, prominence. (E) Same as (D) but at PCW 9.5. Insets show the meningeal clusters, and where scRNA-seq was sampled. Colours for meningeal layers match those in (C). CNS, central nervous system; ECA, epicranial aponeurosis; LM, leptomeninges; SMC, smooth muscle cells. (F) Analysis of meningeal layer maturation. Top left: UMAP of all fibroblasts, subsets of PCW 6-, 10-, and 13- (this study), and adult^11^ fibroblasts. Cells are coloured by their expression of the gene they express most highly among *LAMC3* (pia), *SLC22A6* (arachnoid), *COL8A1* (dura). Bottom left panel: stacked bar chart showing the fraction of fetal and adult fibroblasts expressing a list of enriched genes from adult pia, arachnoid, and dura (Methods). Y, years. Right panel: median expression of the enriched gene lists over developmental timepoints. (G) Top: Immunohistochemistry of ki67 and DAPI at PCW 9.5. Inset region ‘iv’ as in (E), but on an adjacent section. Bottom: scRNA-seq UMAP of fibroblasts coloured by cell cycle score. (H) Spatial transcriptomics showing layer-specific markers. Brackets indicate the meninges. (I) Same as (H) but *SLC47A1* (inner dura) and *MSX2* (outer dura, skull and periosteum). (J) Same region as (H-I) but showing spatial clusters. (K) H&E staining. Black brackets show the meninges, pink brackets the dura. (L) H&E staining of adult human dura.

Next, we charted the distribution of fibroblast cell types *in situ*, in whole heads at PCW 6 & PCW 9.5 (Fig. 4D-E). At PCW 6, early craniofacial fibroblasts or mesenchyme were found in well-defined spatial domains like the mesenchyme of extraocular (eyes), maxillary/mandibular (jaws), otic (future ears), and cranial nerve-associated tissues (Fig. 4D). As mentioned before (Fig. 1), we also identified a spatial cluster representing the earliest meninges described in the literature, the primary meninx, from which we had also obtained scRNA-seq data (panels A-C). The primary meninx was a thin layer surrounding the entire brain together with *FOXQ1*+ endothelial cells (Fig. 4D insets i-ii, Fig. 3I), except around the lower ventral hindbrain where a pool of primary meninx, endothelial-, immune-, and erythropoietic cells were found mixed together (Fig. 4D, inset iii). We found that *LAMC3* was uniquely expressed by the primary meninx (SFig. 4B, Table S4). In contrast, the meningeal marker *FOXC1* had a much broader expression in craniofacial fibroblasts at PCW 6, though it became specific to the meninges at PCW 9.5 (SFig. 4B).

At PCW 9.5, fetal head fibroblasts surrounding the forebrain were already stratified into well-defined layers stacked on top of each other (Fig. 4E, inset iv). Clustering of the spatial data allowed us to identify, from the surface of the head towards the brain, fibroblasts of the skin (periderm, basal epidermis), future skull and periosteum, and the meningeal layers. Other cell types were also present within these fibroblast layers, such as the neural crest-derived Schwann-melanocytic lineage between the periosteum and basal epidermis, lymphatic endothelial cells and pools of erythropoietic cells in the outer periosteum, and classical monocytes mostly in the outer dura. The meninges were divided into eight spatial clusters: five of the outer dura, one inner dura layer, the arachnoid, and the pia.

### The meningeal layers develop concurrently

Next, we focused on meningeal layer formation over time (Fig. 4F-H). Using the scRNA-seq and spatial transcriptomic data we identified and validated unique markers that can distinguish the pia (*LAMC3*), arachnoid (*SLC22A6*) and dura (*COL8A1*), from PCW 5-13 and in adults (Fig. 4F & H, SFig. 4A). *LAMC3*, which was expressed in the primary meninx at PCW 6, was specifically expressed in the pia in later ages. This indicates that the first meningeal fibroblasts to surround the brain are pia precursors, and/or that the pia develops from the primary meninx. Arachnoid and dura precursors were also present at PCW 5-6, though the emerging spatial organisation of meningeal layers was only found around parts of the hindbrain, in these sections (SFig. 4C). At PCW 9.5, precursors of all three layers covered the entire cortex, and *SLC22A2* which was expressed in more mature arachnoid populations appeared first in the anterior cortex (SFig. 4D).

We wondered if the meningeal layers develop as separate lineages, and if therefore intermediate lineage-specific precursor cell types could be identified, as in haematopoiesis. However, examining the expression of the layer-specific markers *LAMC3*, *SLC22A6* and *COL8A1* in subsets of cells from PCW 6, 10 and 13, instead revealed gene expression gradients and a continuum of pia, arachnoid and dura cells at each age (Fig. 4F top left). The same was true when pooling samples at PCW 5-6, 9-10 and 12-13, ruling out batch effects (SFig. 4E). These observations indicate that the meninges mature as a continuum of cell states, only segregating into distinct layer-specific types at developmental timepoints beyond PCW 13.

To further support this parallel model of meninges development, we created a maturation score based on 50 enriched genes from each adult meningeal layer (Methods) (Fig. 4F bottom left), and examined the temporal emergence of each layer as measured by this score. The analysis revealed that (1) genes expressed by the adult meninges were already expressed in the first trimester, (2) the expression of these genes increased over PCW 5-13, demonstrating a gradual maturation of the meningeal layers, and (3) the pia, arachnoid and dura mature concurrently, not sequentially (Fig. 4F bottom left, and right panel, SFig. 4F). The latter conclusion was also supported by the observation of cycling populations in each layer, by scRNA-seq and Ki67 immunostaining (Fig. 4G). Taken together, these results contradict a branched lineage model as seen in neurogenesis or haematopoiesis.

Instead, they support a model where the meningeal layers are formed concurrently, by a gradual refinement of cell states that only segregate into truly distinct layer identities at later developmental timepoints. This latter model resembles how patterning by morphogenetic gradients and cell-cell interactions influence cell fate and layer stratification^37,38^.

### Identification of an inner and outer dura layer

Next, we investigated the dura in more detail (Fig. 4H-L). Our spatial transcriptomic data identified separate outer and inner layers of the dura (Fig. 4E). While both the outer and inner dura expressed *COL8A1*, the known chondrogenic marker *COL2A1*^13^, and dura marker *FXYD5*^7^, the inner layer was distinguished by high *SLC47A1* expression (previously attributed to the dura border in mice^9^). The outer dura was distinguished from the inner dura by expression of *MSX2*, which was also expressed in osteogenic tissues like the developing skull and periosteum (Fig. 4H-J, SFig. 4G). We identified a spatial cluster representing the potential common progenitor of these cells which joined the outside of the primary meninx to surround the brain, and expressed *MSX2, HHIP,* and *COL2A1* (Fig. 4D ‘Skull/dura progenitors’, SFig.4C, Table S4). This suggests that from PCW 5-6, this lineage, together with the primary meninx, give rise to all the meningeal layers. DAPI fluorescence showed a thick outer dura with sparsely located nuclei. However, H&E staining revealed 8-12 sheets of fibroblasts in the outer dura, each a single cell thick, depositing extracellular matrix (ECM) that becomes compact in the adult dura (Fig. 4K-L). These sheets of ECM-embedded fibroblasts extended all the way to the outermost layer of the head, the periderm, and could represent a general mechanism for fibroblast layer formation.

### The inner dura expresses many tight-junction genes necessary to form a barrier

The inner layer of the dura comprised a sheet of single cells, apposed onto the arachnoid. In the fetal head, the epithelial marker *CDH1* was expressed in three fibroblast layers: the meningeal barrier separating arachnoid and dura, the outermost periosteal layer (or epicranial aponeurosis) and the periderm (SFig. 5A), raising interesting questions about fibroblast organisation and barrier formation during layer development. Focusing on the meningeal barrier, in mice, *Cdh1* is reportedly only expressed by arachnoid barrier cells^39^. However, in our data, *CDH1* was expressed by both the *SLC47A1*+ inner dura and *SLC22A6*+ arachnoid barrier cells, forming two very closely aligned layers of cells (Fig. 5A-B). The inner dura and arachnoid barrier cells and their precursors also shared expression of other genes, such as *CCN3* (also known as *NOV*), *SLC4A4* and *PTGDS* (Fig. 5A-B).

**Figure 5.**
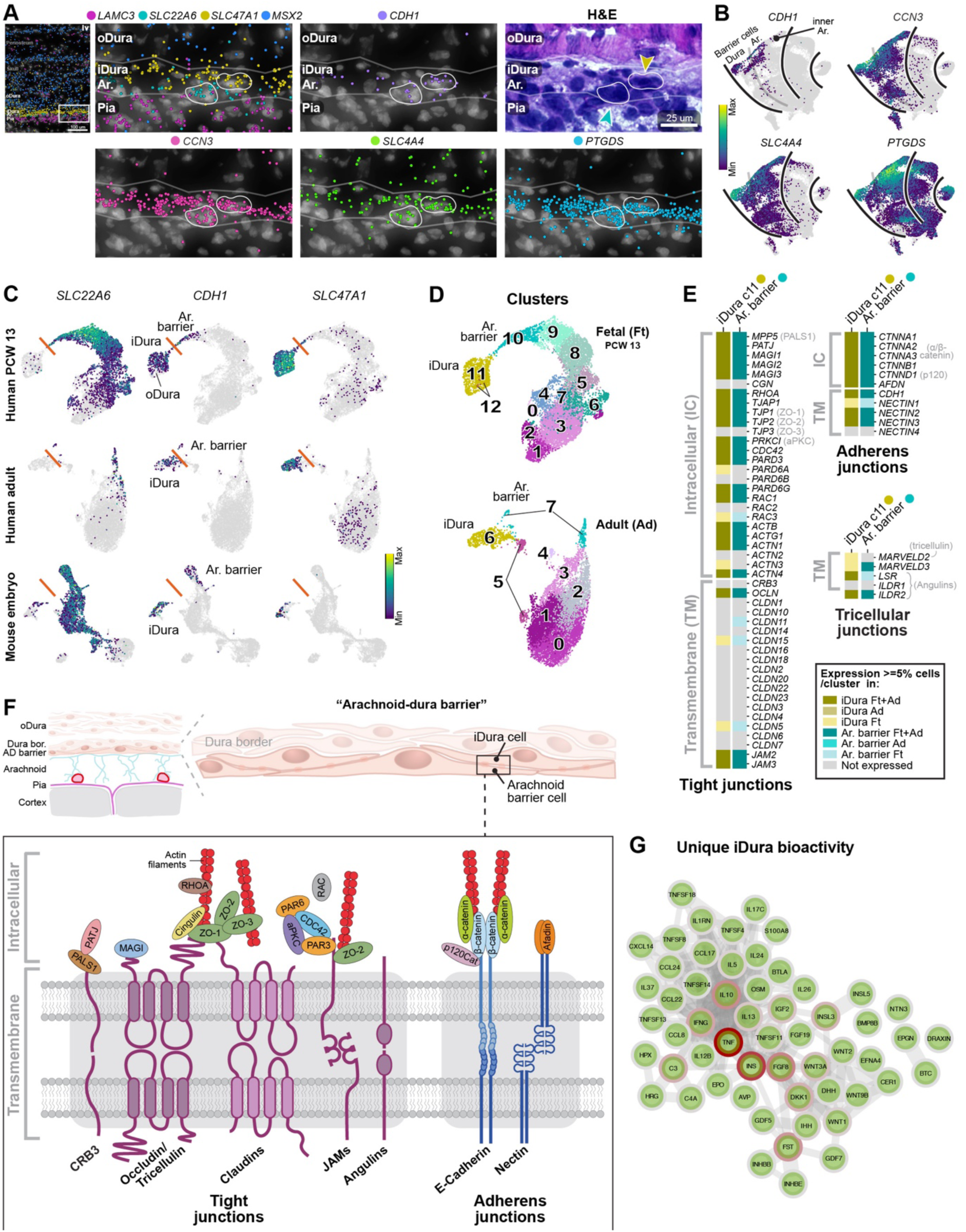
Analysis of the arachnoid barrier and inner dura. (A) Spatial transcriptomics of *CDH1*, *CCN3* (also known as *NOV*), *SLC4A4*, and *PTGDS*, and H&E staining. Ar., arachnoid; iDura, inner dura; oDura, outer dura. (B) Same genes as (A) but coloured on scRNA-seq UMAPs of fibroblasts, on a grey background of all cells. Black lines separate putative pia, arachnoid, and dura lineages. (C) UMAPs with *SLC22A6* (arachnoid), *CDH1* (barrier cells), and *SLC47A1* (inner dura) expression in human fetal (this study), human adult^11^, and mouse embryonic^41^ fibroblasts. Orange lines indicate the separation between arachnoid and dura, based on *SLC22A6* expression. (D) Clusters in fetal PCW 13 fibroblasts (this study) and adult fibroblasts^11^. Cluster colours match the expression of layer-specific genes in (A). (E) Stylised heatmaps showing the expression of genes encoding core components of tight-, adherens- and tricellular junction complexes, in clusters of fetal and adult inner dura-, and arachnoid barrier cells. Grey text indicates protein names where the gene names are dissimilar. (F) Schematic illustration of our hypothesis that the arachnoid barrier is created between a layer of arachnoid barrier cells and the inner dura. Tight- and adherens junction proteins produced by genes in (E) are shown, where the proteins left to right are in the same order as in (E) top to bottom, for intracellular and transmembrane domains. (G) STRING interaction network of a uniquely bioactive gene signature in inner dura. Gene centrality outlined by red.

*PTGDS* has been reported to target the arachnoid^40^, but *Ptgds-Cre* lineage tracing in mice has shown that it is expressed in the whole meninges^5^. In our data, *PTGDS* expression progressively increased with age in all three layers, including the pia, and inside the more developed parts of the future skull (SFig. 5B-C).

These observations suggest that the inner dura layer may form a functionally distinct epithelial layer attached directly on top of the arachnoid, rather than being a simple continuation of the dura proper. Supporting this idea, the inner dura histologically closely resembled the arachnoid barrier rather than the outer dura (Fig. 5A, H&E). Although not previously reported, we noted further that arachnoid and dura cells in both human adult^11^ and mouse fetal meninges^41^ also expressed *CDH1* (Fig. 5C), demonstrating that this inner dura layer is an evolutionarily conserved feature that persists into the adult (Fig. 5C). Spatial transcriptomics of human adult dura demonstrated that there existed *CDH1*+ *SLC47A1*+ cells, and that were negative for *SLC22A6* and *SLC22A2* (SFig. 5D).

To form a seal separating the dural and subarachnoid spaces, arachnoid barrier cells express tight- and adherens junctions. Interestingly, we found that genes encoding many of the components of a functional tight junction complex (Fig. 5D-F) were expressed in both the human fetal and adult inner dura (Fig. 5E, olive colour). Key genes included the transmembrane components *OCLN* (Occludin), a few *CLDN*s (Claudins), and *JAM*s, and the intracellular domains *MAGI*s, *TJP*s (Zona Occludens proteins), *PARD3*/*PARD6A-B* (PAR3/PAR6) and *CDC42*. Likewise, genes encoding the core components of adherens junctions were expressed, like *CDH1* (E-Cadherin), *NECTIN*s, *CTNNA/Bs* (α/β-catenin), and *AFDN* (Afadin) (Fig. 5E-F, SFig. 5E-F). Finally, four components of tricellular junctions were also expressed in the inner dura, *MARVELD2* (Tricellulin), *MARVELD3*, *LSR* (Angulin-1), and *ILDR2* (Angulin-3) which together are sufficient to form functional tricellular junction units (Fig. 5E).

Based on these observations we propose the possibility that the complete meningeal barrier is formed by two *CDH1*+ layers of cells; the inner dura and the arachnoid barrier (Discussion).

We also found signatures of high bioactivity in the developing meningeal barrier (Table S6). STRING network and pathway analysis showed that the inner dura and arachnoid barrier shared the expression of genes involving a strong IL6-central inflammatory/fibrotic network, PI3K/AKT-, WNT-, and VEGFA/VEGFR2-signaling, developmental biology and axon guidance signatures, EGF-IL6-WNT5A-central cancer pathways, and an IL1B-FN1-central immune network (SFig. 5G). However, inner dura cells uniquely expressed a network of bioactive genes with a strong cytokine and growth factor signature. TNF and INS were the most central genes, followed by FGF8, IL10, FST, C3, and IFNG (Fig. 5G).

Looking at enriched genes in the inner dura and arachnoid barrier, ontologies of shared enriched genes included ‘Cell junction’ and ‘Anchoring junction’ (Table S6). Ontologies of uniquely enriched genes in the inner dura represented ‘Extracellular matrix’ and ‘Secreted’, while the arachnoid barrier had an enrichment of ‘Transport of small molecules’ and ‘Active transmembrane transporter activity’ via solute carriers (SFig. 5H). Therefore, the function of the meningeal barrier may also be questioned if the cellular composition includes inner dura cells.

### Meningioma tumours are dura-like

Meningioma tumours are thought to arise from the meningothelial cells of the arachnoid, encompassing both arachnoid cap and barrier cells, based on their histological similarities^3^. However, the discovery of inner dura cells calls into question the precise cell of origin of meningiomas: is it arachnoid meningothelial cells or the inner dura? Like most meningiomas, both cell types expressed *CDH1*. Both also expressed *PTGDS*, and thus cannot be distinguished based on findings in mice, where meningioma can be generated by expression of *NF2* in *PTGDS*-expressing primordial meningeal cells^40^. To address the question, we therefore reanalysed previously published scRNA-seq data^18^, and generated new spatial transcriptomic data from seven meningiomas. We compared tumour data to our fetal scRNA-seq and spatial data, and to previously published adult meninges snRNA-seq^11^ and bulk^18^ datasets.

We processed ten sections from seven meningiomas (six hypermitotic grade 3, one grade 1, Table S1) with spatial transcriptomics using the same gene panel as the fetal samples (Table S3). To allow comparison with fetal meninges, we applied latent Dirichlet allocation (LDA) topic modeling^42^ to our scRNA-seq data from PCW 5-13 fibroblasts and perivascular cells (Fig. 6A, SFig. 6A-B). Like clustering, LDA identifies topics — gene modules — that correspond to cell identities. In addition, it can reveal modules that are shared across cell types, such as cell cycle genes or genes involved in transcriptional reactive states. We generated 35 topics and identified those corresponding to the pia, arachnoid, *PTGDS*+ precursors, inner dura and dura (inner + outer) (Fig. 6A). We also identified one topic shared between arachnoid and inner dura cells; topic #20, ‘barrier’, with the junctional gene *CDH1* appearing as one of the topic-defining genes (SFig. 6C, Table S7). To validate these inferred topic identities, we transferred the topics to fetal spatial transcriptomic data. In agreement, we found that they were highly active in the corresponding anatomical layers of the meninges. As expected, the barrier topic #20 was shared; i.e. active in both arachnoid and inner dura layers (Fig. 6B-C).

**Figure 6.**
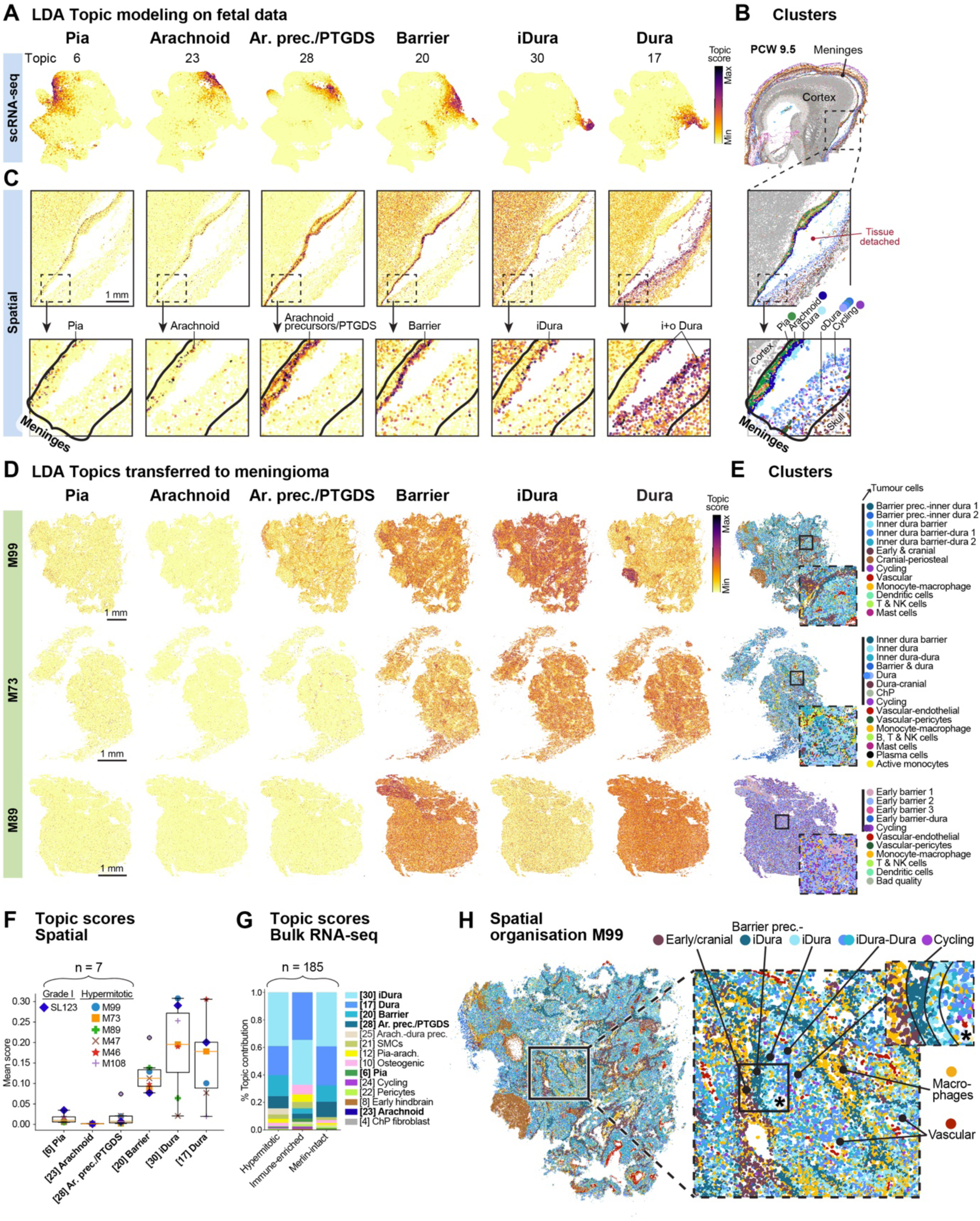
LDA topic modeling in fetal meninges and meningioma tumours. (A) UMAP of fetal fibroblasts and perivascular cells, coloured by their LDA topic score, with topics annotated. Ar. prec., arachnoid precursor; iDura, inner dura; oDura, outer dura. (B) PCW 9.5 spatial clusters. Insets highlight clusters of meningeal layers. (C) LDA topics transferred to fetal spatial data. Insets are the same as in (B). (D) LDA topics transferred to spatial data of grade 3 hypermitotic meningiomas M99, M73 & M89. (E) Spatial clusters of tumours. Clusters were coloured by their most similar meningeal layer (e.g. inner dura-like cells were coloured cyan, as in (B)). (F) Boxplots showing scores from LDA topic transfer to spatial data from seven meningiomas. (G) Stacked bar chart showing scores of LDA topics transferred to bulk RNA-sequencing data from 185 meningiomas, separated into three methylation profiles. (H) Insets of M99 spatial clusters.

Having thus confirmed that topic modelling identified gene programs specific to meningeal layers, we next transferred the topics to tumour spatial data (Fig. 6D-F, SFig. 6D). Although tumour cells are highly abnormal, we reasoned that the expression of normal gene programs might still reveal their underlying cell identity. As expected, we found that the pia topic was nearly absent. More surprisingly, the arachnoid topic was also absent; in fact, it was the least active topic in every tumour examined.

Instead, the most highly activated topics were the inner dura, dura and barrier. Notably, in normal meninges, the barrier topic was shared between arachnoid and inner dura cells, but the absence of arachnoid topic activity in meningiomas argues strongly that tumour cells were not arachnoid. Instead, barrier topic activity in tumour cells coincided with inner dura or dura topics only. Considering the presence of gene programs related to some meningeal layers, this also informed the annotation of meningiomas (Fig. 6E, SFig. 6E). Furthermore, looking at the expression of standard layer markers we did not notice any *SLC22A6* expression (arachnoid), while *CDH1* (barrier) and *SLC47A1* (inner dura) were clearly expressed. Consistently, where *PTGDS* was expressed, it was in non-arachnoid cells (SFig. 6F).

Extending the analysis to a larger cohort of tumours, we transferred our topics to a previously published bulk RNA-seq dataset of 185 meningiomas (hypermitotic, immune-enriched and Merlin-intact subtypes; Fig. 6G, SFig. 6G)^18^. In agreement with the findings from our spatial data, these bulk datasets showed very low activity of pia and arachnoid topics. Instead, inner dura and dura topics dominated, followed by the barrier topic in hypermitotic and merlin-intact meningiomas, and an osteogenic topic in immune-enriched meningiomas. Overall, these observations strongly suggest that meningiomas takes on a predominantly dura cell identity, and indicate a dural-lineage cell as the origin of these tumours.

Furthermore, spatial clustering showed that some, but not all, tumours were highly spatially structured in a manner that resembled a jumbled version of dural and early cranial development (Fig 6H).

Tumour cells resembling developmental dural, inner dural and cranial cell types formed nested layers, with vascular and immune cells interspersed. In other tumours, however, cells were mixed more uniformly (Fig. 6E, SFig. 6E). Notably, there were some tumour clusters containing expression of periosteal-, chondrogenic-, and cranial markers (*MFAP5*, *COL2A1*, *HHIP*) (SFig. 6H dotplot, Table S4). These cells had a bone-like appearance with H&E staining (SFig. 6H). Cycling cells, vasculature and macrophages were confirmed with immunohistochemistry, and we also observed that tumours contained lectin-positive non-vascular regions (SFig. 6H). Finally, we collected four human adult dura samples for spatial transcriptomics. The dura samples were obtained from patients with either low-grade glioma or subependymoma, i.e. non-invasive tumours where the meninges should be unaffected (’normal’). While cycling tumour cells expressed *CDH1*, *CCN3*, and *SLC47A1*, and were therefore inner dura-like, the only cycling cells in normal adult dura were vascular, and not dural fibroblasts (SFig. 6H).

### The dura lineage specifically expresses genes frequently mutated in meningioma

Further extending our analysis to a previously analysed scRNA-seq dataset of six meningiomas^18^ (Fig. 7A), we found that again pia and arachnoid topics were nearly absent and instead barrier, inner dura and dura topics dominated in separate clusters (SFig. 7A-D). We were also able to provide additional annotation to this dataset. For example, our ‘Dura-like meningioma’ overlapped the previously annotated ‘ECM remodelling meningioma’, and a subset of pericytes were ‘pia-like’ (Fig. 7B, SFig. 7A-D). Furthermore, applying an improved karyotyper algorithm (Methods) to this scRNA-seq dataset we found that most cycling cells were derived from tumour MSC6, which carried a distinct karyotype (chromosome 9, 12, and 15 gain) and lacked the typical chromosome 22 loss that was used to define tumour cells in the original publication (Fig. 7C, SFig. 7E).

**Figure 7.**
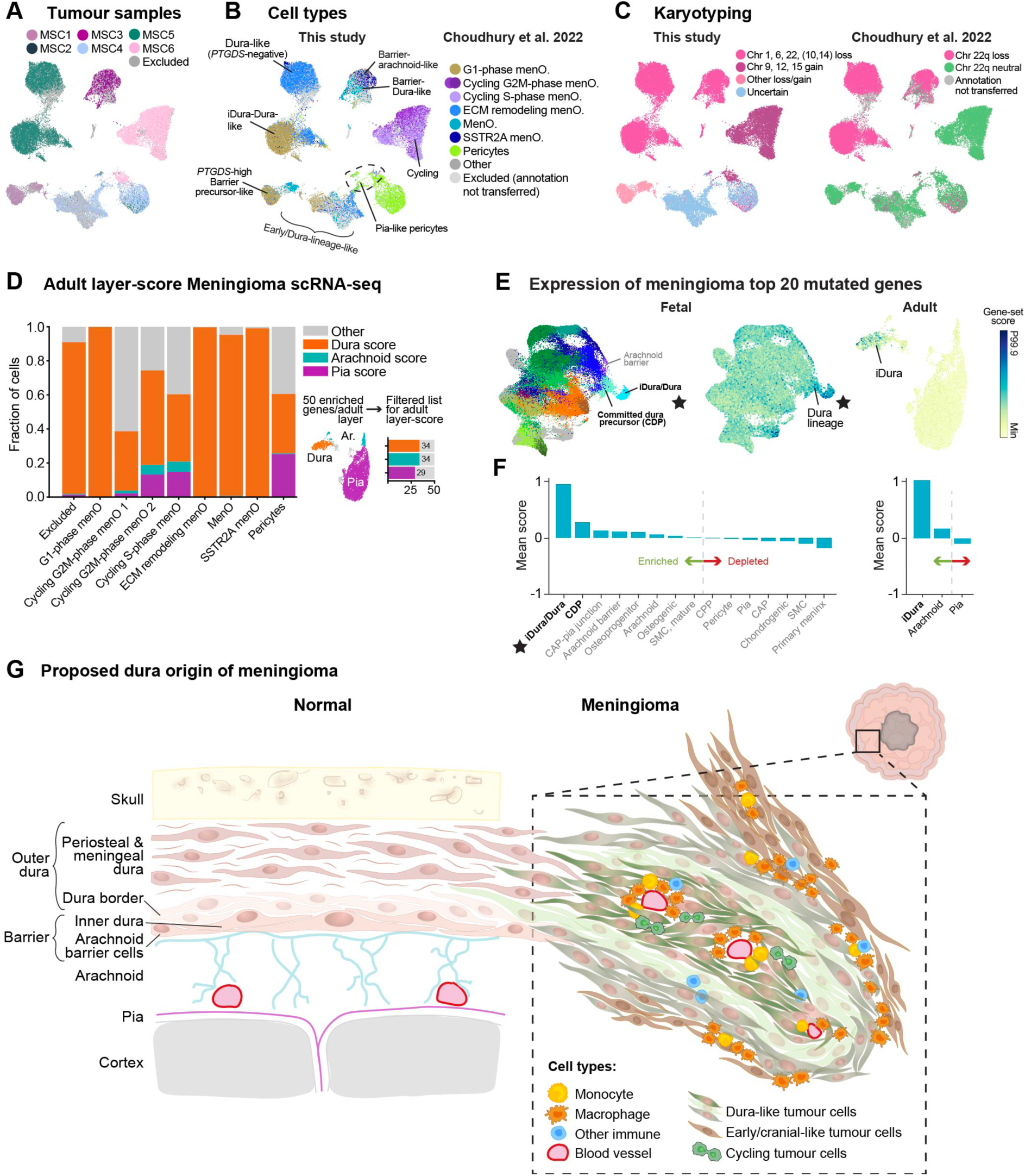
Genetic analysis supporting a dura origin of meningioma tumours. (A) UMAP of tumour cells and pericytes, coloured by samples. Data from Choudhury et al. (2022)^18^. (B) UMAP coloured by transferred annotations^18^, with additional annotations added in this study. Arach., arachnoid; ECM, extracellular matrix; iDura, inner dura; MenO, meningioma. (C) UMAP coloured by karyotype. (D) Stacked bar chart showing the fraction of tumour cell types expressing enriched genes from adult meningeal layers (as in Figure 4). Lane for immune cells not shown in Figure, since we focused on tumour cells. (E) Left: UMAP of fetal fibroblasts and perivascular cells, coloured by Subclass. Right: UMAPs of fetal fibroblasts and perivascular cells and adult fibroblasts^11^, coloured by a gene-set score of the top 20 mutated genes in meningioma (https://cancer.sanger.ac.uk/cosmic/browse/tissue, Table S8). (F) Mutated genes-score as in (E) quantified per fetal Subclass. Stars indicate the same cells across (E-F). CDP, committed dura precursor. (G) Schematic illustration of healthy meninges, and the proposed formation of meningioma tumours from dura cells.

The LDA topics above were defined on developing meninges, but tumours are found in postnatal brains. As an alternative indicator of cell identity, we therefore next identified layer-specific genes expressed preferentially in adult meninges (exactly as previously described for Figure 4). Examining their expression in tumour cells, we found that in each subtype of tumour cells, the dura score dominated, while arachnoid was never among the top three. Only the pia score approached that of dura, and only in pericytes (Fig. 7D). These observations provide strong evidence in favour of a dural cell identity in meningiomas.

Gene expression can provide evidence for the cell of origin, but it remains possible that the fully developed tumour has taken on an identity different from the cell of origin. As an independent line of evidence based on cancer genomics, we reasoned that most cancer driver genes must be normally expressed in the cell of origin — or how else would they transform that cell? We therefore collected the 20 most frequently mutated genes in meningioma (COSMIC Cancer Browser, SFig. 7H) and analysed their collective expression in adult and fetal meninges. Intriguingly, the cell type with by far the highest expression of these putative meningioma cancer drivers in the developing meninges was the inner dura-dura, followed by committed dura precursors. (Fig. 7E-F, left). In healthy adult meninges, putative meningioma driver genes were expressed preferentially in inner dura cells (Fig. 7E-F, right). To reduce the influence of common cancer driver genes shared with other cancer types, we repeated the analysis after removing genes commonly mutated also in glioblastoma, intestinal cancer or in pancreatic cancer (Table S8). In each case, inner dura-dura and/or committed dura precursors remained the most enriched for expression of cancer drivers (SFig. 7G-L). These results lend further independent support to dural cells as the origin of meningioma, and particularly singles out inner dura cells as the most likely candidate. We therefore propose a model whereby meningioma tumours arise from cells of the dural lineage (Fig. 7G).

## DISCUSSION

Fibroblasts and mesenchymal cells are poorly described in general, partly due to their unconventional lineage plasticity^43,44^. This often leads to ambiguous descriptions of cellular subtypes and function, and poor subtype markers (e.g. the canonical marker *COL1A1* is expressed in fibroblasts, all perivascular cells, and osteochondral cells). Spatial transcriptomics greatly aided a more detailed and accurate annotation of fetal craniofacial and meningeal fibroblasts. By using scRNA-seq and spatial transcriptomics we molecularly described the earliest progenitors of pia, arachnoid and dura mater. We also showed that meningeal layer development is defined by gene expression gradients, and occurs concurrently (Fig. 4).

### Hypothesis of inner dura cells as a second barrier sheet on top of the arachnoid

Importantly, we discovered a fibroblast type transcriptionally similar to dura cells (*FXYD5*, *SLC47A1*) but expressing *CDH1*; i.e. ‘inner dura’ cells. In the fetal meninges, these cells formed a single sheet directly on top of *CDH1*+ arachnoid barrier precursor cells (Fig. 5). The current literature reports that only arachnoid barrier cells express *CDH1*^39^, and a scRNA-seq atlas of the adult human leptomeninges annotated *CDH1*+*PTGDS*+ cells as the arachnoid barrier^10^. The first scRNA-seq paper of mouse E14.5 *Col1a1*+ cells found two clusters expressing *Cdh1*, one arachnoid (cluster ‘M3’) and one dura (‘M4’). However, it was reported that only the ‘M3’ arachnoid cluster expressed tight junction genes^7^. Interestingly, our inner dura cells expressed many genes encoding a functional tight junction, and looked more like arachnoid barrier than outer dura cells histologically. This raises the intriguing possibility that the inner dura layer is epithelial (*CDH1*), and that the complete meningeal barrier is formed by two layers of cells; the inner dura and the arachnoid barrier. This model is further supported by electron microscopy^9^, which has shown the meningeal barrier to be formed by precisely two layers of cells. It would also agree with the fact that in our tissue samples, the inner dura cells remained attached to the arachnoid (not the rest of the dura) at sites where the meninges had partially ruptured. Our findings are not necessarily inconsistent with mouse models of arachnoid barrier layer development^45^, but show that the use of *CDH1* (and *PTGDS*) does not discriminate between the arachnoid barrier and inner dura, and therefore propose that the composition of the ‘arachnoid barrier’ may include the inner dura. If future experiments confirm this hypothesis, we propose renaming the arachnoid barrier to the ‘arachnoid-dura barrier’, and the *CDH1*+ inner dura cells to ‘dura barrier cells’.

### Hypothesis of meningioma cell of origin as dura cells, probably inner dura

Meningiomas have been described based on their location and histological subtypes, and classified based on atypical features like mitosis, and invasion^3^. However, they don’t always behave according to their grade, in terms of unexpected recurrences and malignant transformations^46^. Recent efforts have refined meningioma classifications based on integrative multidimensional molecular profiling, and paired them with clinical parameters to more accurately reflect clinical outcomes^47–50^. Surprisingly though, meningiomas have not been described based on their molecular similarity to meningeal layers. Potentially, this has been due to the longstanding acceptance of the meningothelial arachnoid cell origin^51,52^, and the existence of very few meningeal molecular and cellular atlases^7–10^ to compare with. Here, we leveraged our comprehensive atlas of human meninges development to show that meningiomas contain cells similar to cells of the dura lineage. Instead, arachnoid topics were depleted (Fig. 6). Evidence supporting more specifically inner dura cells as a meningioma progenitor cell, was that they (1) expressed *CDH1*, as did meningioma, (2) were located in the meningeal barrier, consistent with previous anatomical evidence, (3) histologically looked like arachnoid barrier cells, (4) of all the topics, inner dura cells (topic 30) were the most consistent in all meningiomas (Fig. 6), and (5) expressed cancer driver genes. The fact that genetic driver genes of meningioma were enriched in inner dura cells (Fig. 7) suggests that these cells are particularly vulnerable to cancer transformation.

Previous studies were able to generate meningiomas from *PTGDS*-expressing primordial meningeal cells in mouse-models^4,53^. However, we demonstrated that *PTGDS* doesn’t distinguish the arachnoid barrier from the inner dura. While our conclusion contradicts the current view of arachnoid meningothelial cells as the cell of origin for meningiomas, our findings are not necessarily inconsistent with previous findings, just that they did not discriminate between arachnoid barrier and inner dura cells experimentally and histologically. Furthermore, considering the clinical diversity of meningioma tumours it is likely that other progenitor cells drive or contribute to meningioma tumorigenesis as well, such as, for example, *NOTCH3*+ mural cells^5^.

### Limitations of the study

Fetal tissue availability was a limiting factor, for example, we miss PCW 11 meninges (Fig. 1). The majority of dura cells in our scRNA-seq atlas expressed *CDH1* (Fig. 5), therefore, we captured mostly inner dura. Likewise, we aspired to collect the arachnoid and whole dura from adult patients for spatial analysis, but due to surgical limitations we mostly obtained the dura border to some periosteal dura.

To confirm barrier function of the inner dura, the expression of tight junction proteins need to be confirmed, together with functional experimentation. Hence, we presented the results of our analysis as a hypothesis for the inner dura forming a barrier with the arachnoid barrier cells.

In this study, we mainly focused on hypermitotic grade 3 meningiomas for spatial transcriptomics (Fig. 6). However, meningiomas have 15 histological subtypes^54^. While we performed LDA topic transfer to bulk RNA-seq of 185 meningiomas (grades 1-3), further investigation is necessary to conclude whether our observations apply to other meningioma subtypes, or clinical parameters (age, sex, survival etc.).

## RESOURCE AVAILABILITY

### Lead contact

Further information and requests for resources and reagents should be directed to and will be fulfilled by the lead contact, Sten Linnarsson.

### Materials availability

This study did not generate new unique reagents.

### Data and code availability

- scRNA-seq: BAM files are available from the European Genome/Phenome Archive (https://ega-archive.org/) under accession number TBA. Count matrices are available from our companion GitHub page at https://github.com/linnarsson-lab/human-meninges-development.
- Xenium *in situ* data and images are available from the BioImage Archive under accession number S-BIAD1600 (https://www.ebi.ac.uk/bioimage-archive/).
- All original code for the analysis and visualisation of data has been deposited to our companion GitHub page at https://github.com/linnarsson-lab/human-meninges-development.
- All of the above are publicly available as of the date of publication.
- Any additional information required to reanalyse the data reported in this paper is available from the lead contact upon request.

## ACKNOWLEDGMENTS

Firstly, we thank the donors for the samples used in this study. We thank Abrar Choudhury for transfer of meningioma metadata, and Kimberly Siletti for the initial implementation of LDA topic modeling. We thank Lars Borm for freezing and sharing the fetal heads. We thank Natalie Welsh, and Ezequiel Goldschmidt, for their insight about tight junctions and comparing the developing meninges and meningioma tumours respectively. We thank members of the Linnarsson lab for helpful discussions and lab management. We acknowledge Katarina Tiklova and the In Situ Sequencing Facility at SciLifeLab, funded by Science for Life Laboratory and the Swedish Research Council, for providing in situ sequencing services. We acknowledge support from the National Genomics Infrastructure in Stockholm, and the Biomedicum Imaging Core facility.

## AUTHOR CONTRIBUTIONS

Conceptualisation, E.V., and S.L.; scRNA sequencing, L.H., and E.V.; scRNA data preprocessing, P.L.; scRNA data analysis, E.V., P.L., and S.L.; B lineage analysis and visualisation, S.W.; immunohistochemistry, I.K., and E.V.; Xenium data processing, S.M.S.; Xenium data analysis, E.V., and S.M.S.; data annotation, E.V.; visualisation, E.V.; data curation and software, P.L., and S.M.S.; resources, X.H., R.A.B., E.S., X.L., D.R.R., M.N., M.H., and O.P.; writing - original draft, E.V., S.L., and S.W.; funding acquisition, S.L., M.N., and E.V.

All authors read and approved the final version of the manuscript.

## SUPPLEMENTAL INFORMATION

**Document S1. Figures S1-S7**

**Table S1. Samples**

**Table S2. scRNA-seq cluster metadata**

**Table S3. Xenium probe panels**

**Table S4. Xenium clusters**

**Table S5. CSF analysis**, related to Figure S3

**Table S6. Network analysis**, related to Figures 5 and S5

**Table S7. Topics filtered genes**, related to Figures 6, S6, and S7

**Table S8. COSMIC mutated genes**, related to Figure 7 and S7

## METHODS

### KEY RESOURCES TABLE

| REAGENT or RESOURCE | SOURCE | IDENTIFIER |
| --- | --- | --- |
| **Antibodies** |  |  |
| Recombinant Anti-Ki67 antibody [B56] | Abcam | Cat# ab279653, RRID: AB_2934265 |
| Aquaporin 1/AQP1 Antibody (1/22) | Santa Cruz Biotechnology | Cat# sc-32737, RRID: AB_626693 |
| Anti-Iba1 antibody [EPR16588] | Abcam | Cat# ab178846, RRID: AB_2636859 |
| Anti-alpha smooth muscle Actin antibody [SP171] | Abcam | Cat# ab150301 |
| Goat anti-Mouse IgG (H+L) Highly Cross-Adsorbed Secondary Antibody, Alexa Fluor™ Plus 555 | Thermo Fisher Scientific | Cat# A32727, RRID: AB_2633276 |
| Goat anti-Rabbit IgG (H+L) Highly Cross-Adsorbed Secondary Antibody, Alexa Fluor™ 647 | Thermo Fisher Scientific | Cat# A-21245, RRID: AB_2535813 |
| Goat anti-Mouse IgG2b Cross-Adsorbed Secondary Antibody, Alexa Fluor™ 555 | Thermo Fisher Scientific | Cat# A-21147, RRID: AB_2535783 |
| **Biological samples** |  |  |
| Fetal meninges | Karolinska Hospital (Danderyd & Huddinge), Addenbrooke's Hospital Cambridge | N/A |
| Fetal heads | Addenbrooke's Hospital Cambridge | N/A |
| Human adult dura | Karolinska Hospital (Department of neurosurgery) | N/A |
| Human meningioma | University of California San Francisco | N/A |
| **Chemicals, peptides, and recombinant proteins** |  |  |
| Sodium hydrosulfite | Sigma-Aldrich | Cat# 71699 |
| 10× PBS, pH 7.4, RNase-free | Thermo Fisher Scientific | Cat# AM9625 |
| Hematoxylin and Eosin Stain Kit | Vector laboratories | Cat# H-3502 |
| Xylenes | Sigma-Aldrich | Cat# 534056 |
| Entellan™ new | Sigma-Aldrich | Cat# 1.07961 |
| Pierce™ 16% Formaldehyde (w/v), Methanol-free | Thermo Fisher Scientific | Cat# 28908 |
| Pierce™ 20X TBS Buffer | Thermo Fisher Scientific | Cat# 28358 |
| Triton™ X-100 | Sigma-Aldrich | Cat# T8787 |
| Normal goat serum | Thermo Fisher Scientific | Cat# 31873 |
| BSA (30%) | Sigma-Aldrich | Cat# A9576 |
| Hoechst 33258 solution | Sigma-Aldrich | Cat# 94403 |
| Lectin HPA From *Helix pomatia* (edible snail), Alexa Fluor™ 488 Conjugate | Thermo Fisher Scientific | Cat# L11271 |
| ProLong™ Gold Antifade Mountant | Thermo Fisher Scientific | Cat# P36930 |
| Hibernate™-E Medium | Thermo Fisher Scientific | Cat# A1247601 |
| 2-Methylbutane | Sigma-Aldrich | Cat# M32631 |
| Isopropanol | Sigma-Aldrich | Cat# 34863 |
| Nuclease-free dH2O | Thermo Fisher Scientific | Cat# 10977035 |
| Papain Dissociation System | Worthington | Cat# LK003150 |
| **Critical commercial assays** |  |  |
| Chromium Single Cell 3' GEM, Library & Gel Bead Kit v3, 16 rxns | 10x Genomics | Cat# PN-1000075 |
| Chromium Single Cell B Chip Kit, 48 rxns | 10x Genomics | Cat# PN-1000073 |
| Chromium i7 Multiplex Kit, 96 rxns | 10x Genomics | Cat# PN-120262 |
| Xenium Slides & Sample Prep Reagents | 10X Genomics | Cat# 1000460 |
| Xenium Decoding Reagents | 10X Genomics | Cat# 1000461 |
| Xenium Decoding Consumables | 10X Genomics | Cat# 1000487 |
| Xenium Human Lung Gene Expression Panel | 10X Genomics | Cat# 1000601 |
| **Deposited data** |  |  |
| scRNA-seq, human fetal meninges | This Manuscript | https://cellxgene.cziscience.com/collections/7d66d871-091f-4602-9f42-85f86d2853e0 |
| Xenium in situ, human fetal heads | This Manuscript | https://www.ebi.ac.uk/bioimage-archive/ , #S-BIAD1600 |
| Xenium in situ, human meningioma | This Manuscript | https://www.ebi.ac.uk/bioimage-archive/ , #S-BIAD1600 |
| Xenium in situ, human dura | This Manuscript | https://www.ebi.ac.uk/bioimage-archive/ , #S-BIAD1600 |
| Xenium & scRNA-seq loom files | This Manuscript | https://github.com/linnarsson-lab/human-meninges-development?tab=readme-ov-file |
| Human fetal endothelial cells | Braun et al.<sup>12</sup> | doi: 10.1126/science.adf1226 |
| Human adult fibroblasts | Siletti et al.<sup>11</sup> | doi: 10.1126/science.add7046 |
| Mouse embryonic fibroblasts | La Manno et al.<sup>41</sup> | doi: 10.1038/s41586-021-03775-x |
| Meningioma scRNA-seq, bulk RNA-seq | Choudhury et al.<sup>18</sup> | doi: 10.1038/s41588-022-01061-8 |
| CSF proteomics human adult | Guldbrandsen et al.<sup>35</sup>, Dayon et al.<sup>36</sup> | doi: 10.1074/mcp.M114.038554, doi: 10.1021/acs.jproteome.8b00809 |
| **Oligonucleotides** |  |  |
| Human custom probe panel | 10X Genomics | Table S3 |
| Human Lung Gene Expression Panel | 10X Genomics | https://www.10xgenomics.com/products/xenium-panels |
| **Software and algorithms** |  |  |
| Cytograph | Siletti et al.<sup>11</sup> | https://github.com/linnarsson-lab/adult-human-brain/tree/main/cytograph |
| Souporcell | Heaton et al.<sup>19</sup> | https://github.com/wheaton5/souporcell |
| Tomotopy v0.12.2 | Blei et al.<sup>42</sup> | https://bab2min.github.io/tomotopy/v0.13.0/en/ |
| STAR v2.7.10a | Dobin et al.<sup>55</sup> | https://github.com/alexdobin/STAR |
| CellRanger v6.1.2 | 10X Genomics | https://www.10xgenomics.com/support/software/cell-ranger/latest |
| Cytoscape v3.10.1 | Shannon et al.<sup>56</sup> | https://cytoscape.org |
| STRING v2.2.0 | Szklarczyk et al.<sup>57</sup> | https://www.string-db.org |
| Xenium data processing | Salas et al.<sup>58</sup>, this manuscript | https://www.biorxiv.org/content/10.1101/2023.02.13.528102v1 |
| Code for analysis and visualisation | This manuscript | https://github.com/linnarsson-lab/human-meninges-development |
| **Other** |  |  |
| Superfrost Plus™ Adhesion Microscope Slides | Epredia | Cat# J1800ABDH |
| ImmEdge Hydrophobic Barrier PAP Pen | Vector laboratories | Cat# H-4000 |
| Microscope Cover Glasses No. 1, 24 × 50mm | Nordic biolabs | Cat# 101222 |
| 30 µm cell strainer (CellTrics) | Sysmex | Cat# 04-0042-2316 |
| Cancer genes | COSMIC | https://cancer.sanger.ac.uk/cosmic/browse/tissue |
| Xenium in situ platform | 10X Genomics | N/A |

## EXPERIMENTAL MODEL AND STUDY PARTICIPANT DETAILS

### Donors

Human prenatal samples were collected from elective medical abortions at the Department of Gynecology, Danderyd Hospital and Karolinska Huddinge Hospital, and Addenbrooke’s Hospital in Cambridge, following oral and written informed consent by the patient. In Sweden, the use of abortion material was approved by the Swedish Ethical Review Authority and the National Board of Health and Welfare. In the UK, approval from the National Research Ethics Service Committee East of England – Cambridge Central was obtained (Local Research Ethics Committee, 96/085).

#### Patient samples

Human dura samples and one meningioma sample (“SL123”) were collected from the Karolinska Hospital with informed consent from the patients and with ethical approval from the Swedish Ethical Review Authority (2020-03505). The use of samples was approved by the Swedish Ethical Review Authority (2020-02096).

Meningioma samples were collected from the University of California San Francisco (UCSF), complied with all relevant ethical regulations, and was approved by the UCSF Institutional Review Board (13–12587, 17–22324, 17–23196 and 18–24633). As part of routine clinical practice at UCSF, all patients who were included in this study signed a written waiver of informed consent to contribute deidentified data to research projects.

Details about all samples can be found in Table S1.

## METHOD DETAILS

### Tissue sample collection

#### Human fetal meninges

14 samples of human prenatal meninges were used in this study, at post-conception weeks (PCW) 5-13. Two samples of fetal heads were used, at PCW 6 and 9.5. The post-conception age of the embryos and fetuses was estimated by information from the clinical ultrasound, last menstrual period, true crown-rump-length, and age-dependent anatomical landmarks.

For samples collected at the Karolinska Hospital, the tissue was immediately transported to the laboratory following abortion, and dissected in ice-cold 0.9% NaCl solution under sterile conditions within 1-2 hours post abortion. For scRNA-seq, meninges were dissected and kept in ice-cold Hibernate-E medium until further processing. For spatial analysis fetal heads were covered by Tissue-Tek Optimal Cutting Temperature compound (OCT) in cryomolds, snap-frozen in a slurry of 2-methylbutane (Sigma-Aldrich) and dry ice, and stored at −80°C pending sectioning.

For samples collected in Cambridge, tissues were dissected in a class II hood on the day of collection and stored in Hibernate-E medium at 4°C. The tissue was shipped to Sweden at refrigerated temperature, and delivered two days after abortion. The procedure is covered under ethics REC: 96/085.

#### Human adult dura

Four dura samples were collected from patients undergoing surgery for low grade gliomas and subependymoma (i.e. non-invasive, and meninges unaffected). During surgery, dura was placed in Hibernate-E medium on ice, and transported as such. Upon arrival, samples were washed in fresh Hibernate-E, then embedded in OCT, and snap-frozen in 2-methylbutane and dry ice slurry. Samples were stored at −80°C until further use.

#### Human meningioma

Six meningioma tumour samples that were resected from 1991 to 2016 and snap frozen were identified from an institutional biorepository and clinical database at the University of California San Francisco (UCSF), with an emphasis on high-grade meningiomas.

WHO grading was performed using contemporary criteria outlined in the WHO 2016 and 2021 classification of tumours of the central nervous system. Samples were shipped to Sweden on dry ice, and stored at −80°C. Upon use, samples were embedded in OCT for cryosectioning. “SL123” meningioma was collected at the Karolinska Hospital as the dura samples above.

#### Cell dissociation

Dissected meninges were processed around 6 to 48 hours after tissue collection, depending on the source (Karolinska Hospital/Cambridge). Tissues were stored at 4°C in Hibernate-E medium during transport from Cambridge and until processing. Ice cold Earle’s Balanced Salt Solution (Worthington) was carbogenated (95% O2/ 5% CO2) and used throughout the whole procedure. Meninges were dissociated using the Worthington’s Papain Dissociation System (Worthington) (Protocols.io, https://dx.doi.org/10.17504/protocols.io.xmbfk2n). Tissues were enzymatically digested at 37 °C for 10-15 min, followed by trituration using fire polished glass Pasteur pipettes. Cell suspensions were filtered through a 30μm cell strainer (CellTrics, Sysmex) and centrifuged for 5 min at 200 g to obtain cell pellets. Supernatants were carefully removed, and cells resuspended in small volumes of EBSS (depending on cell density). Cell concentrations were estimated using a counting haemocytometer (Bürker/Neubauer chamber) and diluted with EBSS until the desired concentrations were reached. All suspensions were kept on ice until loading on the 10X Chromium chips.

#### Single-cell RNA sequencing

Droplet-based single-cell RNA sequencing was performed using the 10x Genomics Chromium Single Cell Kit v3. Single-cell suspensions concentrated at 800-1200 cells/ml were mixed with master mix and nuclease free water according to the Chromium manual, targeting 5000-10000 cells per reaction. 12 PCR cycles were used for cDNA synthesis, and the rest of the library preparation was performed according to the manufacturer’s instructions (10X Genomics, Illumina). All libraries were sequenced on an Illumina NovaSeq 6000 using S4 to a target sequencing depth of 100,000 reads/cell. Sequencing saturation was examined for each sample using preseq (https://github.com/smithlabcode/preseq).

#### Cryosectioning

Fresh-frozen tissue was embedded in OCT as above. The tissue was acclimatised in the cryostat chamber, and the specimen/chuck was set to −16°C and the knife −14°C (both for fetal heads, dura, and meningiomas). The samples were sectioned at 10μm and collected on Xenium slides following the manufacturer’s recommendations, or captured on Superfrost Plus Adhesion Microscope Slides for other stainings as below.

#### Xenium In Situ Gene Expression

Fresh-frozen 10μm tissue sections were placed on Xenium slides. The next steps were performed at the In Situ Sequencing Facility at SciLifeLab. The tissue was fixed and permeabilized according to the Xenium Fixation and Permeabilization Protocol (Demonstrated Protocol CG000581). A predesigned Human Lung panel (289 genes) and a custom gene panel (100 genes, Table S3) were applied to the tissue. Probes were hybridized to target RNA, followed by ligation and enzymatic amplification to generate multiple copies of each RNA target, as outlined in the Probe Hybridization, Ligation, and Amplification User Guide (User Guide CG000582). The prepared Xenium slides were subsequently loaded onto the Xenium Analyzer for imaging and analysis, following the Decoding and Imaging User Guide (User Guide CG000584). Instrument software version 1.4.3.1 and software analysis version 1.4.0.7 were used throughout the process.

#### Post-Xenium hematoxylin and eosin staining

To remove autofluorescence quencher from the post-Xenium slides, tissues were incubated in 1.74mg/ml sodium hydrosulfite dissolved in Mili-Q water for 10 min. The slides were then rinsed three times in Mili-Q water for 1 min, followed by a 2 min wash with distilled water (dH_2_O). The slides were subsequently washed three times with 1× PBS and three times with dH_2_O for 5 min. Hematoxylin solution (Hematoxylin and eosin stain kit) was applied to each section for 1 min and rinsed in two changes of dH_2_O. Bluing agent (Hematoxylin and eosin stain kit) was applied to each section for 15 seconds and the slide was rinsed twice with dH_2_O. The slide was dipped in ethanol and excess > 99.5% ethanol was blotted off. Eosin Y solution (Hematoxylin and eosin stain kit) was applied to each section for 30 seconds and rinsed in ethanol. The sections were subsequently dehydrated in three changes of > 99.5% ethanol for 2 min each and rinsed in xylenes, before mounting with Entellan™ new. The slide was left to solidify for 24 hours in a ventilated hood.

#### Immunohistochemistry cryosections

Fresh-frozen tissue samples were cryosectioned into 10μm sections and captured on Superfrost Plus Adhesion Microscope Slides. A hydrophobic barrier was drawn around tissue sections using ImmEdge Hydrophobic Barrier PAP Pen. The sections were fixed with 4% formaldehyde in 1× PBS for 10 min. The slides were subsequently washed with 1× TBS for 5 min, then washed with three changes of 0.025% Triton X-100 in 1× TBS for 5 min each with gentle agitation. The sections were incubated with 200 µl of 10% normal goat serum and 1% bovine serum albumin (BSA) in 1× TBS blocking solution per section for 2 hours at room temperature. A mixture of desired primary antibodies was prepared with following dilutions: 1:1000 Recombinant Anti-Ki67 antibody [B56] and either 1:100 Aquaporin 1/AQP1 Antibody (1/22), 1:1000 Anti-Iba1 antibody [EPR16588], or 1:200 Anti-alpha smooth muscle Actin antibody [SP171] with 1% BSA in 1× TBS. The sections were incubated with the primary antibody mixture in a humidifying chamber at 4°C overnight. The slides were washed with two changes of 0.025% Triton X-100 in 1× TBS for 5 min each with gentle agitation. A mixture of secondary antibodies was prepared with following dilutions: 1:500 Goat anti-Mouse IgG (H+L) Highly Cross-Adsorbed Secondary Antibody Alexa Fluor™ Plus 555 (for use with Ki67 primary antibody) and either 1:500 Goat anti-Rabbit IgG (H+L) Highly Cross-Adsorbed Secondary Antibody Alexa Fluor™ 647 (for use with IBA1 or ACTA2 primary antibodies) or 1:500 Goat anti-Mouse IgG2b Cross-Adsorbed Secondary Antibody, Alexa Fluor™ 555 (for use with AQP1 primary antibody) with 1% BSA in 1× TBS. The sections were incubated with the secondary antibody mixture for 1.5 hours at room temperature in a dark staining chamber. Subsequently, the slide was washed with three changes of 1× TBS for 5 min each in a dark humidifying chamber. A stain mixture was prepared containing 2 μg/ml Hoechst 33258 and 1:200 Lectin HPA From Helix pomatia (edible snail) Alexa Fluor™ 488 Conjugate in 1× TBS. The sections were incubated with the stain mixture for 30 min at room temperature in a dark staining chamber. The slides were washed with 0.025% Triton X-100 in 1 × TBS for 30 min and subsequently washed in three changes of 1× TBS for 5 min each at room temperature in a dark staining chamber. The slides were washed in 1 × TBS for 30 min in a dark staining chamber, before mounting with ProLong™ Gold Antifade Mountant and a coverslip. The slides were left to solidify overnight in a dark staining chamber.

#### Immunohistochemistry free-floating tissue

Fresh meninges pieces were placed into a 12-well plate and fixed with 4% formaldehyde in 1× PBS for 10-15 min with gentle agitation. Subsequent steps were conducted as above but in a 12-well plate, with antibodies diluted in a volume of 1 ml/well to cover the tissue, and washed with 2 ml. Following the final washes, the floating meninges were placed on Superfrost Plus Adhesion Microscope Slides, and excess liquid was carefully removed. They were then mounted with ProLong™ Gold Antifade Mountant and a coverslip as above, and left to solidify overnight in a dark staining chamber

#### Histology image acquisition

Histology image acquisition was performed on a Zeiss Axio Scan.Z1 slide scanner with Zen Blue v3.1 acquisition software, using the Plan-Apochromat 20x/0.8 M27 objective with NA 0.8. For acquiring brightfield images (Hematoxylin and eosin staining) Hitachi HV-F203SCL camera (pixel size of 4.4 µm) with VIS-LED illumination source were used. For acquiring fluorescence images Hamamatsu ORCA-Flash4.0 V3 Digital CMOS camera (pixel size of 6.5 µm) with a Colibri 7 LED light source (100% intensity), PBS 405 + 493 + 575 + 654 + 761 beamsplitter, and PBP 425/30 + 514/31 + 592/25 + 681/45 + 785/38 emission filters. Namely, the following configurations were used: H3258 channel (Hoechst stain) was imaged using LED-Module 385 nm and BP 385/30 excitation filter; AF488 channel (Lectin HPA Alexa Fluor 488 conjugate) was imaged using LED-Module 475nm and BP 469/38 excitation filter; AF555 channel (Goat anti-Mouse IgG Highly Cross-Adsorbed Secondary Antibody, Alexa Fluor Plus 555, Goat anti-Mouse IgG2b Cross-Adsorbed Secondary Antibody, Alexa Fluor 555) was imaged using LED-Module 567 nm and BP 555/30 excitation filter; AF647 channel (Goat anti-Rabbit IgG Highly Cross-Adsorbed Secondary Antibody, Alexa Fluor 647) was imaged using LED-Module 630 nm and BP 631/33. Histology image acquisition was performed at Karolinska Institutet’s Biomedicum Imaging Core facility (https://ki.se/en/bic).

#### Alignment of Xenium and histology images

H&E images are provided in the Xenium data folders (BioImage Archive, accession number S-BIAD1600). To align H&E files to the Xenium data, the correct .ome file, and .csv alignment file needs to be opened in the Xenium browser.

## QUANTIFICATION AND STATISTICAL ANALYSIS

### scRNA-seq data preprocessing

Illumina runs were demultiplexed with cellranger mkfastq version 6.1.2 (10x Genomics). Read mapping and unique molecular identifier (UMI) counts were determined using STARSolo^55^ version 2.7.10a, using human genome GRCh38.p12 and transcript annotations from ENSEMBL release 93 from reference package GRCh38-3.0.0 as available from 10Xgenomics (www.10xgenomics.com). STARSolo was run with the following parameters:

--soloType CB_UMI_Simple

--soloCellFilter EmptyDrops_CR <expectedNcells> 0.99 10 45000 90000 500 0.01 20000 0.01 10000

--soloCBmatchWLtype 1MM_multi_Nbase_pseudocounts

--soloUMIfiltering MultiGeneUMI_CR

--soloUMIdedup 1MM_CR

--clipAdapterType CellRanger4

--outFilterScoreMin 30

Replicates were pooled, resulting in one loom file (loompy.org) per sample.

#### Quality control

Samples were analysed with “cytograph qc” (https://github.com/linnarsson-lab/adult-human-brain), which uses a modified version of DoubletFinder to calculate a doublet score for each cell^59^. We captured a median 13062 cells from each sample and then filtered cells based on their total number of mRNA molecules—as counted by unique molecular identifiers (UMIs)—and percentages of unspliced RNA, as well as doublet scores. Cytograph qc was initially run with standard parameters, where cells with fewer than 1000 UMIs, unspliced molecule fraction less than 0.1, or a doublet score below 0.4 were removed from further analysis. A second round of QC was performed after clustering (below).

After QC, a final dataset containing 156,726 cells remained. Mean cells, genes and UMIs per sample are in Table S1. Variability across donors likely reflected the quality of the tissue.

#### Clustering

Cells were clustered using an updated version of the Cytograph package (https://github.com/ linnarsson-lab/adult-human-brain). All cells that passed QC were pooled into a single dataset for initial clustering. The command “cytograph build” was run with configuration factorization: HPF, nn_space: HPF, n_factors: 94, steps: nn, embeddings, clustering, aggregate, export. Default values were used for other parameters.

To get finer clusters, these top-level clusters were split by the dendrogram using the command “cytograph split --method dendrogram”. Fibroblasts, neural, and vascular cells were split one more time by the dendrogram such that no split had more than 37 clusters. A second round of more stringent quality control was performed, whereby a cluster was removed if at least 40% of cells in the cluster had fewer than 1800 UMIs, or if we manually spotted clusters containing doublets or *HBB* (erythrocyte) contamination. Then, the splits were pooled into final subsets; Subset_Neural, Subset_Neural_crest, Subset_Epithelial, Subset_Fibroblasts, Subset_Perivascular, Subset_Endothelial, Subset_Immune, Subset_Erythropoietic (Table S2). Subset clusters were annotated and analysed separately, and also pooled into a final complete dataset containing all 245 clusters (Table S2; All_cells). UMAP embeddings were calculated for each subset, and the pooled dataset of all cells.

#### Annotation

Clusters were manually annotated as CellType based on literature, regional, and age composition (Table S2). Three additional levels of annotations were provided, on broader levels; “Subclass”, “Class”, and “Superclass”. We computed cell cycle scores as previously described^41^. We used the expression of a set of well-known cell cycle genes^60^ as a proxy for active proliferation. We calculated the cell cycle score as the fraction of UMIs those genes represented, and then the percentage of cells per cluster with a cell cycle score > 0.01. We also computed gene enrichment as previously described^41^, and available in deposited Jupyter Notebooks (https://github.com/linnarsson-lab/human-meninges-development). Briefly, gene enrichment is a measure of overexpression in a cluster relative to other clusters, taking into account both mean expression and fraction of non-zero cells.

#### Maternal calling by SNP analysis

We used Souporcell as created by Heaton et al., 2020^19^, and freely available at https://github.com/wheaton5/souporcell, applying it to the output from the STARSolo runs. Souporcell uses variants detected in scRNA-seq reads to assign cells to their donor of origin. In order to make sure that we applied Souporcell correctly, we first ran it on a dataset collecting separate maternal and fetal samples (data not shown) before analysing our own dataset.

### Processing of Xenium’s spatial datasets

Four types of samples were profiled using Xenium: (1) developmental human meninges (PCW 6), (2) developmental human meninges (PCW 9.5), (2) adult human dura and (3) meningioma samples. Due to the differences expected between each of the sample types, each of the sample types were processed independently using Scanpy (Code availability).

#### Processing of developmental and adult meningeal Xenium datasets

For developmental (PCW 6 and PCW 9.5) heads and adult dura samples, cells were first loaded and combined into a single AnnData object. With the aim of minimising the segmentation errors caused by Xenium’s segmentation pipeline, only transcripts identified within segmented nuclei were kept for each of the profiled cells. This approach discards a considerable amount of the profiled reads but minimises potential mis-segmentation effects (Marco Salas et al., 2023^58^). Only transcripts annotated as genes in the provided panel were retained. Cells were then filtered to remove low-quality observations, discarding cells with less than eight detected genes or fewer than 20 total transcripts. After this, expression values were normalised to a total count of 10,000 transcripts per cell, followed by logarithmic transformation. Highly variable genes were identified based on specific thresholds of mean expression and dispersion, and principal component analysis (PCA) was performed to reduce dimensionality. The neighbourhood graph was constructed using the top 50 principal components, and clustering was conducted using the Leiden algorithm. A UMAP 2D representation was generated to visualise cell clusters, and the processed data were saved. Based on the spatial location and differentially expressed genes identified for every cluster, clusters were annotated assigning a cell type to each high-quality cell profiled. For developmental datasets (PCW 6 and PCW 9.5), due to the complexity of both mesenchymal and neural clusters, these clusters were independently subclustered in a subsequent analysis (Code availability), integrating the resulting annotations into a final object containing a cell type assigned to each high-quality cell profiled.

#### Processing of meningioma Xenium datasets

Due to the high biological variability between tumour samples, meningioma’s processing was done on a tumour-basis, combining only replicates of the same tumour when available. In here, quality control measures were also applied to filter out cells with fewer than three detected genes or fewer than five total transcripts. Expression data were normalised to a total count of 10,000 transcripts per cell, followed by a logarithmic transformation. Principal component analysis (PCA) was used to reduce dimensionality, with the top 50 principal components selected to capture major sources of variation in the dataset. A neighbourhood graph was constructed using the top 40 components, enabling (1) the application of the Leiden algorithm to identify clusters and (2) a low-dimensional representation of the data, using UMAP.

#### B lineage analysis

We produced dotplots to visualise gene expression, using raw gene expression count matrices subset to cell types of interest as input. We then normalised these count matrices using the Scanpy (v1.9.3) sc.pp.normalise_per_cell function with counts_per_cell_after=1e4. A ln(x+1) transformation was then applied to the normalised counts using the sc.pp.log1p function. Gene expression was then visualised using the sc.pl.dotplot function. To increase interpretability of plots, the vmax argument was sometimes used to apply an upper limit to the color scale (where used, noted within GitHub scripts).

Data produced in this study was compared to data from fetal bone marrow as B-cell reference (Jardine et al., 2021)^22^, and yolk sac (Goh et al., 2023)^21^.

#### Neftel score

We independently reimplemented the gene profile scoring algorithm from Neftel et al., 2019^61^, and generalised it to work with any given set of genes. Our implementation in Python is provided as neftel_score.py, and we confirmed that our code reproduces the original analysis from Neftel et al.

#### Fibroblast layer-maturation score

Fibroblasts from the snRNA-seq data from Siletti et al., 2023^11^ was downloaded, and top 50 enriched genes per meningeal layer calculated. Three criteria were imposed to filter the gene lists such that they contained genes that are highly expressed, and after PCW 6.

1. Criteria to be considered expressed in PCW 5-6: >15% of cells should have at least one transcript of the gene. If expressed in PCW 5-6, these genes were removed so that the gene list reflects genes involved in maturation of the meningeal layers.
2. Criteria to be considered expressed: Gene count above 2.
3. Criteria to be considered pia/arachnoid/dura: At least 5% of the enriched genes should be expressed in a cell to not become “Other”.

#### STRING Network and pathways analysis

The gene expression values of a list of bioactive molecules, and enriched genes, for the inner dura and arachnoid barrier clusters was analysed and visualised using Cytoscape version 3.10.1 (Shannon et al., 2003^56^). The expression of bioactive molecules and enriched genes were analysed (1) as uniquely expressed by the inner dura or arachnoid barrier, and (2) as shared between the two.

STRING database^57^ version 2.2.0 was used to visualise networks of interconnected molecules (50% confidence for node connectivity), belonging to significantly enriched pathway ontologies (Table S6). Pathway analysis in STRING is based on KEGG, Reactome, and Wiki Pathways.

#### Topic modeling with Latent Dirichlet Allocation (LDA)

LDA^42^ was used to identify transcriptional modules that vary across cells, and also reveals co-expressed genes in single-cell data. We used tomotopy version 0.12.2 to perform topic modeling with LDA. Model training, identification of topic-representative genes, and other general parameters were performed as previously described^11^. Here, the number of topics “k” generated were 35. Some topics were specific to quality, for example *HBB* erythrocyte contamination, mitochondrial or ribosomal genes, and one topic contained genes from the Y-chromosome. Some topics were not specific or did not seem to be biologically meaningful. Therefore, we manually chose topics of interest. We also identified representative genes for each topic by using the topic probabilities reported for each gene (function “get_topic_word_dist”). We filtered genes by specificity (topic probabilities for each gene normalized by that gene’s probability across all topics) and then sorted the remaining genes by their unnormalized probabilities (Table S7). A script is available as optimize_lda.py at github.com/linnarsson-lab/adult-human-brain, and scripts of its application in this study are available at https://github.com/linnarsson-lab/human-meninges-development.

LDA also allows you to score new cells for a given gene topic. Hence, we validated the scRNA-seq topic annotations by transfer to spatial data (scoring spatial data for the LDA topics generated on scRNA-seq data).

#### Transfer of scRNA-seq LDA topics to Xenium in situ gene expression data

Topics from the model were transferred to matching genes in the Xenium dataset and the weights obtained used to colorize the *in situ* images.

#### Transfer of scRNA-seq LDA topics to bulk RNA-seq data

Bulk RNA-seq data of 185 meningiomas was downloaded from Choudhury et al. (2022)^18^ and split according to their methylation group; Merlin-intact, Immune-enriched, and Hypermitotic. For each tumor and tumor type, the relative contribution of each topic was calculated, and sums were normalized to 1.0.

### Reanalysis of meningioma scRNA-seq data

Downloaded from Choudhury et al. (2022)^18^ and processed with the same Cytograph pipeline as described for the fetal meninges scRNA-seq data generated in this study. Tumour cells (fibroblast-like) and pericytes were isolated into a new loom file, to keep the comparison consistent with the fetal meningeal fibroblasts and pericytes.

### Computational expression-based karyotyping

We inferred chromosomal copy number aberrations by determining normalized gene expression along chromosomes, using a normal brain atlas (Siletti et al. 2023)^11^ as the reference. By using an external reference, we avoided the need to manually identify normal cells within each sample. We normalized expression in each cell to the median total UMIs across all cells. We included only genes from the autosomes that were expressed at more than 10% of the 99^th^ percentile expression level in at least half of the clusters in the reference atlas. Then, for each cell in the sample, we identified the reference cell type (cluster in the atlas) with the greatest correlation coefficient. For each cell, we then computed a ploidy score along the chromosomes by dividing the normalized expression in the cell by the normalized expression in the reference cell type and multiplying by two. We smoothed the ploidy along the chromosomes in windows of 25 genes and then normalized the resulting ploidy to the median ploidy and multiplied by two. We used the resulting ploidy vectors (one for each cell in the sample) as the input for a clustering procedure: we computed the first five principal components, calculated a nearest-neighbour graph (k=30 neighbours) and used Leiden graph clustering to find metacells having similar ploidy profiles. Finally, for each metacell we used a hidden Markov model to infer integer chromosomal copy numbers. The detailed Python code implementing this algorithm is provided as hmm_karyotyper.py, which also provides visualization of the result.

### Expression of COSMIC mutated genes in fetal meningeal fibroblasts

The Catalogue of Somatic Mutations in Cancer (COSMIC) Cancer Browser (https://cancer.sanger.ac.uk/cosmic/browse/tissue) version 98-99 was used to find the top 20 mutated genes in meningioma, glioblastoma (Astrocytoma Grade IV), intestinal carcinoma, and prostate adenocarcinoma. Overlapping genes between cancers were removed, to ensure that generic cancer drivers didn’t have an effect (Table S8).

## REFERENCES

1. Aldinger, K.A., Lehmann, O.J., Hudgins, L., Chizhikov, V.V., Bassuk, A.G., Ades, L.C., Krantz, I.D., Dobyns, W.B., and Millen, K.J. (2009). FOXC1 is required for normal cerebellar development and is a major contributor to chromosome 6p25.3 Dandy-Walker malformation. Nat. Genet. 41, 1037–1042.

2. Brastianos, P.K., Galanis, E., Butowski, N., Chan, J.W., Dunn, I.F., Goldbrunner, R., Herold-Mende, C., Ippen, F.M., Mawrin, C., McDermott, M.W., et al. (2019). Advances in multidisciplinary therapy for meningiomas. Neuro. Oncol. 21, i18–i31.

3. WHO Classification of Tumours Editorial Board (2021). Central Nervous System Tumours 5th ed. D. N. Louis, ed. (International Agency for Research on Cancer).

4. Kalamarides, M., Stemmer-Rachamimov, A.O., Niwa-Kawakita, M., Chareyre, F., Taranchon, E., Han, Z.-Y., Martinelli, C., Lusis, E.A., Hegedus, B., Gutmann, D.H., et al. (2011). Identification of a progenitor cell of origin capable of generating diverse meningioma histological subtypes. Oncogene 30, 2333–2344.

5. Choudhury, A., Cady, M.A., Lucas, C.-H.G., Najem, H., Phillips, J.J., Palikuqi, B., Zakimi, N., Joseph, T., Birrueta, J.O., Chen, W.C., et al. (2024). Perivascular NOTCH3+ stem cells drive meningioma tumorigenesis and resistance to radiotherapy. Cancer Discov. 14, 1823–1837.

6. Gold, M.P., Ong, W., Masteller, A.M., Ghasemi, D.R., Galindo, J.A., Park, N.R., Huynh, N.C., Donde, A., Pister, V., Saurez, R.A., et al. (2024). Developmental basis of SHH medulloblastoma heterogeneity. Nat. Commun. 15, 270.

7. DeSisto, J., O’Rourke, R., Jones, H.E., Pawlikowski, B., Malek, A.D., Bonney, S., Guimiot, F., Jones, K.L., and Siegenthaler, J.A. (2020). Single-cell transcriptomic analyses of the developing meninges reveal meningeal fibroblast diversity and function. Dev. Cell 54, 43–59.e4.

8. Wang, A.Z., Bowman-Kirigin, J.A., Desai, R., Kang, L.-I., Patel, P.R., Patel, B., Khan, S.M., Bender, D., Marlin, M.C., Liu, J., et al. (2022). Single-cell profiling of human dura and meningioma reveals cellular meningeal landscape and insights into meningioma immune response. Genome Med. 14, 49.

9. Pietilä, R., Del Gaudio, F., He, L., Vázquez-Liébanas, E., Vanlandewijck, M., Muhl, L., Mocci, G., Bjørnholm, K.D., Lindblad, C., Fletcher-Sandersjöö, A., et al. (2023). Molecular anatomy of adult mouse leptomeninges. Neuron 111, 3745–3764.e7.

10. Kearns, N.A., Iatrou, A., Flood, D.J., De Tissera, S., Mullaney, Z.M., Xu, J., Gaiteri, C., Bennett, D.A., and Wang, Y. (2023). Dissecting the human leptomeninges at single-cell resolution. Nat. Commun. 14, 7036.

11. Siletti, K., Hodge, R., Mossi Albiach, A., Lee, K.W., Ding, S.-L., Hu, L., Lönnerberg, P., Bakken, T., Casper, T., Clark, M., et al. (2023). Transcriptomic diversity of cell types across the adult human brain. Science 382, eadd7046.

12. Braun, E., Danan-Gotthold, M., Borm, L.E., Lee, K.W., Vinsland, E., Lönnerberg, P., Hu, L., Li, X., He, X., Andrusivová, Ž., et al. (2023). Comprehensive cell atlas of the first-trimester developing human brain. Science 382, eadf1226.

13. Farmer, D.T., Mlcochova, H., Zhou, Y., Koelling, N., Wang, G., Ashley, N., Bugacov, H., Chen, H.-J., Parvez, R., Tseng, K.-C., et al. (2021). The developing mouse coronal suture at single-cell resolution. Nat. Commun. 12, 4797.

14. Li, B., Li, J., Fan, Y., Zhao, Z., Li, L., Okano, H., and Ouchi, T. (2023). Dissecting calvarial bones and sutures at single-cell resolution. Biol. Rev. Camb. Philos. Soc. 98, 1749–1767.

15. Sankowski, R., Süß, P., Benkendorff, A., Böttcher, C., Fernandez-Zapata, C., Chhatbar, C., Cahueau, J., Monaco, G., Gasull, A.D., Khavaran, A., et al. (2024). Multiomic spatial landscape of innate immune cells at human central nervous system borders. Nat. Med. 30, 186–198.

16. Mortus, J.R., Zhang, Y., and Hughes, D.P.M. (2014). Developmental pathways hijacked by osteosarcoma. Adv. Exp. Med. Biol. 804, 93–118.

17. Okano, A., Miyawaki, S., Hongo, H., Dofuku, S., Teranishi, Y., Mitsui, J., Tanaka, M., Shin, M., Nakatomi, H., and Saito, N. (2021). Associations of pathological diagnosis and genetic abnormalities in meningiomas with the embryological origins of the meninges. Sci. Rep. 11, 6987.

18. Choudhury, A., Magill, S.T., Eaton, C.D., Prager, B.C., Chen, W.C., Cady, M.A., Seo, K., Lucas, C.-H.G., Casey-Clyde, T.J., Vasudevan, H.N., et al. (2022). Meningioma DNA methylation groups identify biological drivers and therapeutic vulnerabilities. Nat. Genet. 54, 649–659.

19. Heaton, H., Talman, A.M., Knights, A., Imaz, M., Gaffney, D.J., Durbin, R., Hemberg, M., and Lawniczak, M.K.N. (2020). Souporcell: robust clustering of single-cell RNA-seq data by genotype without reference genotypes. Nat. Methods 17, 615–620.

20. O’Byrne, S., Elliott, N., Rice, S., Buck, G., Fordham, N., Garnett, C., Godfrey, L., Crump, N.T., Wright, G., Inglott, S., et al. (2019). Discovery of a CD10-negative B-progenitor in human fetal life identifies unique ontogeny-related developmental programs. Blood 134, 1059–1071.

21. Goh, I., Botting, R.A., Rose, A., Webb, S., Engelbert, J., Gitton, Y., Stephenson, E., Quiroga Londoño, M., Mather, M., Mende, N., et al. (2023). Yolk sac cell atlas reveals multiorgan functions during human early development. Science 381, eadd7564.

22. Jardine, L., Webb, S., Goh, I., Quiroga Londoño, M., Reynolds, G., Mather, M., Olabi, B., Stephenson, E., Botting, R.A., Horsfall, D., et al. (2021). Blood and immune development in human fetal bone marrow and Down syndrome. Nature 598, 327–331.

23. Popescu, D.-M., Botting, R.A., Stephenson, E., Green, K., Webb, S., Jardine, L., Calderbank, E.F., Polanski, K., Goh, I., Efremova, M., et al. (2019). Decoding human fetal liver haematopoiesis. Nature 574, 365–371.

24. Vanlandewijck, M., He, L., Mäe, M.A., Andrae, J., Ando, K., Del Gaudio, F., Nahar, K., Lebouvier, T., Laviña, B., Gouveia, L., et al. (2018). A molecular atlas of cell types and zonation in the brain vasculature. Nature 554, 475–480.

25. Kalucka, J., de Rooij, L.P.M.H., Goveia, J., Rohlenova, K., Dumas, S.J., Meta, E., Conchinha, N.V., Taverna, F., Teuwen, L.-A., Veys, K., et al. (2020). Single-cell transcriptome atlas of Murine endothelial cells. Cell 180, 764–779.e20.

26. Cao, J., O’Day, D.R., Pliner, H.A., Kingsley, P.D., Deng, M., Daza, R.M., Zager, M.A., Aldinger, K.A., Blecher-Gonen, R., Zhang, F., et al. (2020). A human cell atlas of fetal gene expression. Science 370, eaba7721.

27. Winkler, E.A., Kim, C.N., Ross, J.M., Garcia, J.H., Gil, E., Oh, I., Chen, L.Q., Wu, D., Catapano, J.S., Raygor, K., et al. (2022). A single-cell atlas of the normal and malformed human brain vasculature. Science 375, eabi7377.

28. Hupe, M., Li, M.X., Kneitz, S., Davydova, D., Yokota, C., Kele, J., Hot, B., Stenman, J.M., and Gessler, M. (2017). Gene expression profiles of brain endothelial cells during embryonic development at bulk and single-cell levels. Sci. Signal. 10. 10.1126/scisignal.aag2476.

29. Jain, A., Ang, P.S., Matrongolo, M.J., and Tischfield, M.A. (2023). Understanding the development, pathogenesis, and injury response of meningeal lymphatic networks through the use of animal models. Cell. Mol. Life Sci. 80, 332.

30. Wilting, J., Papoutsi, M., Christ, B., Nicolaides, K.H., von Kaisenberg, C.S., Borges, J., Stark, G.B., Alitalo, K., Tomarev, S.I., Niemeyer, C., et al. (2002). The transcription factor Prox1 is a marker for lymphatic endothelial cells in normal and diseased human tissues. FASEB J. 16, 1271–1273.

31. Wigle, J.T., Harvey, N., Detmar, M., Lagutina, I., Grosveld, G., Gunn, M.D., Jackson, D.G., and Oliver, G. (2002). An essential role for Prox1 in the induction of the lymphatic endothelial cell phenotype. EMBO J. 21, 1505–1513.

32. Jung, E., Gardner, D., Choi, D., Park, E., Jin Seong, Y., Yang, S., Castorena-Gonzalez, J., Louveau, A., Zhou, Z., Lee, G.K., et al. (2017). Development and characterization of A novel Prox1-EGFP lymphatic and Schlemm’s canal reporter rat. Sci. Rep. 7. 10.1038/s41598-017-06031-3.

33. Antila, S., Karaman, S., Nurmi, H., Airavaara, M., Voutilainen, M.H., Mathivet, T., Chilov, D., Li, Z., Koppinen, T., Park, J.-H., et al. (2017). Development and plasticity of meningeal lymphatic vessels. J. Exp. Med. 214, 3645–3667.

34. Dani, N., Herbst, R.H., McCabe, C., Green, G.S., Kaiser, K., Head, J.P., Cui, J., Shipley, F.B., Jang, A., Dionne, D., et al. (2021). A cellular and spatial map of the choroid plexus across brain ventricles and ages. Cell 184, 3056–3074.e21.

35. Guldbrandsen, A., Vethe, H., Farag, Y., Oveland, E., Garberg, H., Berle, M., Myhr, K.-M., Opsahl, J.A., Barsnes, H., and Berven, F.S. (2014). In-depth characterization of the cerebrospinal fluid (CSF) proteome displayed through the CSF proteome resource (CSF-PR). Mol. Cell. Proteomics 13, 3152–3163.

36. Dayon, L., Cominetti, O., Wojcik, J., Galindo, A.N., Oikonomidi, A., Henry, H., Migliavacca, E., Kussmann, M., Bowman, G.L., and Popp, J. (2019). Proteomes of paired human cerebrospinal fluid and plasma: Relation to blood-brain barrier permeability in older adults. J. Proteome Res. 18, 1162–1174.

37. Mosby, L.S., Bowen, A.E., and Hadjivasiliou, Z. (2024). Morphogens in the evolution of size, shape and patterning. Development 151. 10.1242/dev.202412.

38. Zhu, X.-J., Liu, Y., Dai, Z.-M., Zhang, X., Yang, X., Li, Y., Qiu, M., Fu, J., Hsu, W., Chen, Y., et al. (2014). BMP-FGF signaling axis mediates Wnt-induced epidermal stratification in developing mammalian skin. PLoS Genet. 10, e1004687.

39. Betsholtz, C., Engelhardt, B., Koh, G.Y., McDonald, D.M., Proulx, S.T., and Siegenthaler, J. (2024). Advances and controversies in meningeal biology. Nat. Neurosci. 27, 2056–2072.

40. Peyre, M., Salaud, C., Clermont-Taranchon, E., Niwa-Kawakita, M., Goutagny, S., Mawrin, C., Giovannini, M., and Kalamarides, M. (2015). PDGF activation in PGDS-positive arachnoid cells induces meningioma formation in mice promoting tumor progression in combination with Nf2 and Cdkn2ab loss. Oncotarget 6, 32713–32722.

41. La Manno, G., Siletti, K., Furlan, A., Gyllborg, D., Vinsland, E., Mossi Albiach, A., Mattsson Langseth, C., Khven, I., Lederer, A.R., Dratva, L.M., et al. (2021). Molecular architecture of the developing mouse brain. Nature 596, 92–96.

42. Blei, D.M., Ng, A.Y., and Jordan, M.I. (2003). Latent Dirichlet Allocation. J. Mach. Learn. Res. 3, 993–1022.

43. LeBleu, V.S., and Neilson, E.G. (2020). Origin and functional heterogeneity of fibroblasts. FASEB J. 34, 3519–3536.

44. Plikus, M.V., Wang, X., Sinha, S., Forte, E., Thompson, S.M., Herzog, E.L., Driskell, R.R., Rosenthal, N., Biernaskie, J., and Horsley, V. (2021). Fibroblasts: Origins, definitions, and functions in health and disease. Cell 184, 3852–3872.

45. Derk, J., Como, C.N., Jones, H.E., Joyce, L.R., Kim, S., Spencer, B.L., Bonney, S., O’Rourke, R., Pawlikowski, B., Doran, K.S., et al. (2023). Formation and function of the meningeal arachnoid barrier around the developing mouse brain. Dev. Cell 58, 635–644.e4.

46. Goldbrunner, R., Minniti, G., Preusser, M., Jenkinson, M.D., Sallabanda, K., Houdart, E., von Deimling, A., Stavrinou, P., Lefranc, F., Lund-Johansen, M., et al. (2016). EANO guidelines for the diagnosis and treatment of meningiomas. Lancet Oncol. 17, e383–91.

47. Nassiri, F., Liu, J., Patil, V., Mamatjan, Y., Wang, J.Z., Hugh-White, R., Macklin, A.M., Khan, S., Singh, O., Karimi, S., et al. (2021). A clinically applicable integrative molecular classification of meningiomas. Nature 597, 119–125.

48. Driver, J., Hoffman, S.E., Tavakol, S., Woodward, E., Maury, E.A., Bhave, V., Greenwald, N.F., Nassiri, F., Aldape, K., Zadeh, G., et al. (2022). A molecularly integrated grade for meningioma. Neuro. Oncol. 24, 796–808.

49. Chen, W.C., Choudhury, A., Youngblood, M.W., Polley, M.-Y.C., Lucas, C.-H.G., Mirchia, K., Maas, S.L.N., Suwala, A.K., Won, M., Bayley, J.C., et al. (2023). Targeted gene expression profiling predicts meningioma outcomes and radiotherapy responses. Nat. Med. 29, 3067–3076.

50. Lucas, C.-H.G., Mirchia, K., Seo, K., Najem, H., Chen, W.C., Zakimi, N., Foster, K., Eaton, C.D., Cady, M.A., Choudhury, A., et al. (2024). Spatial genomic, biochemical and cellular mechanisms underlying meningioma heterogeneity and evolution. Nat. Genet. 56, 1121–1133.

51. Yamashima, T., Kida, S., and Yamamoto, S. (1988). Ultrastructural comparison of arachnoid villi and meningiomas in man. Mod. Pathol. 1, 224–234.

52. Tohma, Y., Yamashima, T., and Yamashita, J. (1992). Immunohistochemical localization of cell adhesion molecule epithelial cadherin in human arachnoid villi and meningiomas. Cancer Res. 52, 1981–1987.

53. Boetto, J., Peyre, M., and Kalamarides, M. (2021). Mouse models in meningioma research: A systematic review. Cancers (Basel) 13, 3712.

54. Louis, D.N., Perry, A., Wesseling, P., Brat, D.J., Cree, I.A., Figarella-Branger, D., Hawkins, C., Ng, H.K., Pfister, S.M., Reifenberger, G., et al. (2021). The 2021 WHO classification of tumors of the Central Nervous System: A summary. Neuro. Oncol. 23, 1231–1251.

55. Dobin, A., Davis, C.A., Schlesinger, F., Drenkow, J., Zaleski, C., Jha, S., Batut, P., Chaisson, M., and Gingeras, T.R. (2013). STAR: ultrafast universal RNA-seq aligner. Bioinformatics 29, 15–21.

56. Shannon, P., Markiel, A., Ozier, O., Baliga, N.S., Wang, J.T., Ramage, D., Amin, N., Schwikowski, B., and Ideker, T. (2003). Cytoscape: a software environment for integrated models of biomolecular interaction networks. Genome Res. 13, 2498–2504.

57. Szklarczyk, D., Kirsch, R., Koutrouli, M., Nastou, K., Mehryary, F., Hachilif, R., Gable, A.L., Fang, T., Doncheva, N.T., Pyysalo, S., et al. (2023). The STRING database in 2023: protein-protein association networks and functional enrichment analyses for any sequenced genome of interest. Nucleic Acids Res. 51, D638–D646.

58. Marco Salas, S., Czarnewski, P., Kuemmerle, L.B., Helgadottir, S., Matsson-Langseth, C., Tismeyer, S., Avenel, C., Rehman, H., Tiklova, K., Andersson, A., et al. (2023). Optimizing Xenium In Situ data utility by quality assessment and best practice analysis workflows. bioRxiv. 10.1101/2023.02.13.528102.

59. McGinnis, C.S., Murrow, L.M., and Gartner, Z.J. (2019). DoubletFinder: Doublet detection in single-cell RNA sequencing data using artificial nearest neighbors. Cell Syst. 8, 329–337.e4.

60. Tirosh, I., Izar, B., Prakadan, S.M., Wadsworth, M.H., 2nd, Treacy, D., Trombetta, J.J., Rotem, A., Rodman, C., Lian, C., Murphy, G., et al. (2016). Dissecting the multicellular ecosystem of metastatic melanoma by single-cell RNA-seq. Science 352, 189–196.

61. Neftel, C., Laffy, J., Filbin, M.G., Hara, T., Shore, M.E., Rahme, G.J., Richman, A.R., Silverbush, D., Shaw, M.L., Hebert, C.M., et al. (2019). An integrative model of cellular states, plasticity, and genetics for glioblastoma. Cell 178, 835–849.e21.

