## Supplementary Information for "Cell atlas of the developing human meninges reveals a dura origin of meningioma"

**The PDF file includes:**

Figs. S1-S7

Table S1-S8 Legends

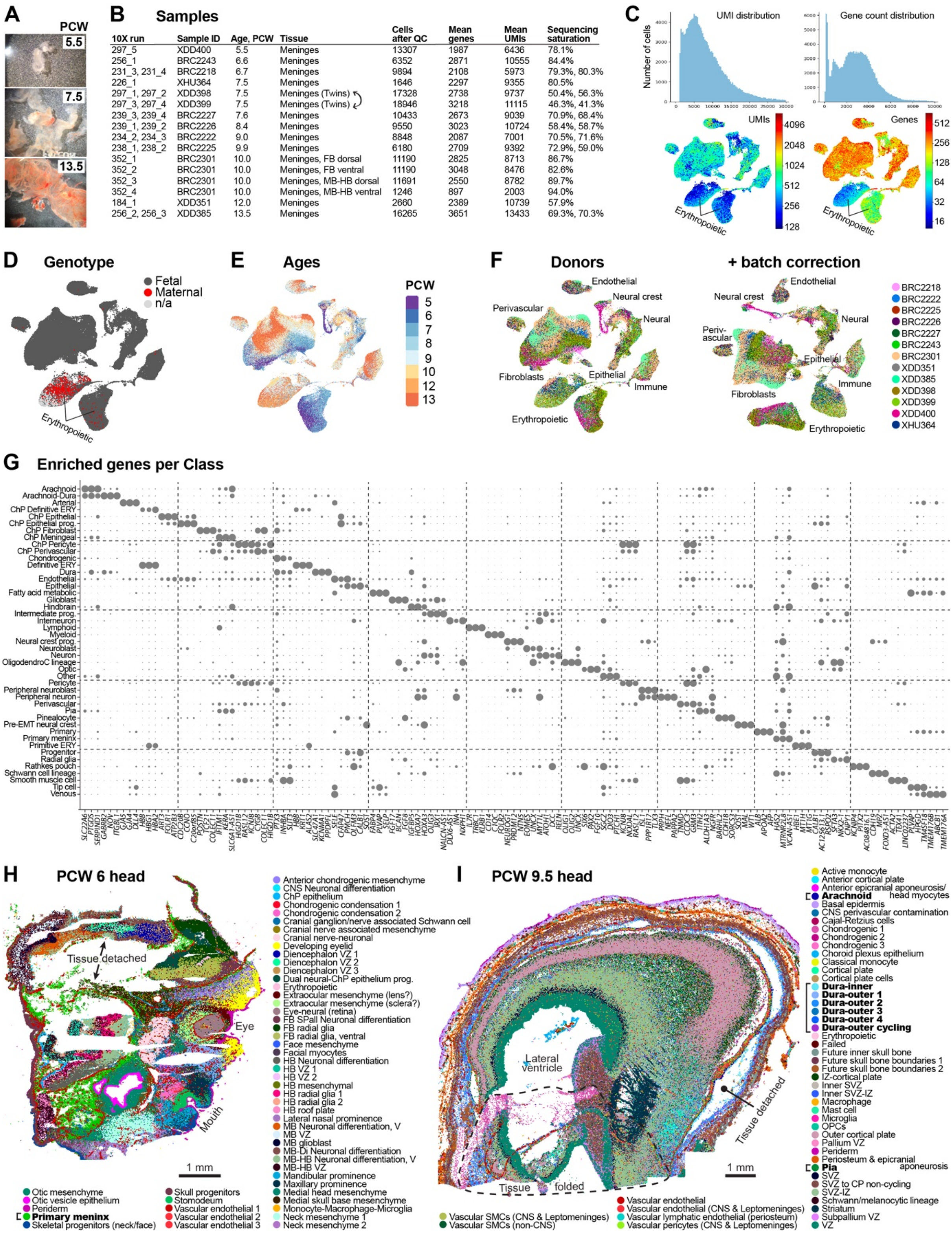


| 10X run | Sample ID | Age, PCW | Tissue | Cells after QC | Mean genes | Mean UMIs | Sequencing saturation |
|---|---|---|---|---|---|---|---|
| 297_5 | XDD400 | 5.5 | Meninges | 13307 | 1987 | 6436 | 78.1% |
| 256_1 | BRC2243 | 6.6 | Meninges | 6352 | 2871 | 10555 | 84.4% |
| 231_3, 231_4 | BRC2218 | 6.7 | Meninges | 9894 | 2108 | 5973 | 79.3%, 80.3% |
| 226_1 | XHU364 | 7.5 | Meninges | 1646 | 2297 | 9355 | 80.5% |
| 297_1, 297_2 | XDD398 | 7.5 | Meninges (Twins) | 17328 | 2738 | 9737 | 50.4%, 56.3% |
| 297_3, 297_4 | XDD399 | 7.5 | Meninges (Twins) | 18946 | 3218 | 11115 | 46.3%, 41.3% |
| 239_3, 239_4 | BRC2227 | 7.6 | Meninges | 10433 | 2673 | 9039 | 70.9%, 68.4% |
| 239_1, 239_2 | BRC2226 | 8.4 | Meninges | 9550 | 3023 | 10724 | 58.4%, 58.7% |
| 234_2, 234_3 | BRC2222 | 9.0 | Meninges | 8848 | 2087 | 7001 | 70.5%, 71.6% |
| 238_1, 238_2 | BRC2225 | 9.9 | Meninges | 6180 | 2709 | 9392 | 72.9%, 59.0% |
| 352_1 | BRC2301 | 10.0 | Meninges, FB dorsal | 11190 | 2825 | 8713 | 86.7% |
| 352_2 | BRC2301 | 10.0 | Meninges, FB ventral | 11190 | 3048 | 8476 | 82.6% |
| 352_3 | BRC2301 | 10.0 | Meninges, MB-HB dorsal | 11691 | 2550 | 8782 | 89.7% |
| 352_4 | BRC2301 | 10.0 | Meninges, MB-HB ventral | 1246 | 897 | 2003 | 94.0% |
| 184_1 | XDD351 | 12.0 | Meninges | 2660 | 2389 | 10739 | 57.9% |
| 256_2, 256_3 | XDD385 | 13.5 | Meninges | 16265 | 3651 | 13433 | 69.3%, 70.3% |

**Figure S1. scRNA-seq QC and spatial clusters, related to Figure 1**
(A) PCW 5.5, 7.5 and 13.5 meninges dissected off fetal brains.
(B) ScRNA-seq samples and sequencing information.
(C) Top: UMI distribution and gene count histograms. Bottom: UMAP embeddings of the scRNA-seq dataset, coloured by log UMIs and log genes per cell.
(D) UMAP coloured by maternal/fetal genotype, from Souporcell SNP analysis.
(E) UMAP coloured by sample ages.
(F) UMAPs without and with Harmony batch correction, coloured by donors.
(G) Dotplot of enriched genes per Class annotation. ChP, Choroid plexus; EMT, epithelial-mesenchymal transition; ERY, erythropoietic; OligodendroC, oligodendrocyte; prog., progenitor.
(H-I) Spatial transcriptomic clusters in sagittal sections of a PCW 6- (H) and PCW 9.5 (I) fetal head. Brackets and bold text highlight meninges-associated clusters. Dashed lines show where tissue folded before analysis. CP, cortical plate; Di, diencephalon; HB, hindbrain; IZ, intermediate zone; MB, midbrain; OPC, oligodendrocyte progenitor cell; SMC, smooth muscle cell; SVZ, subventricular zone; V, ventral; VZ, ventricular zone.

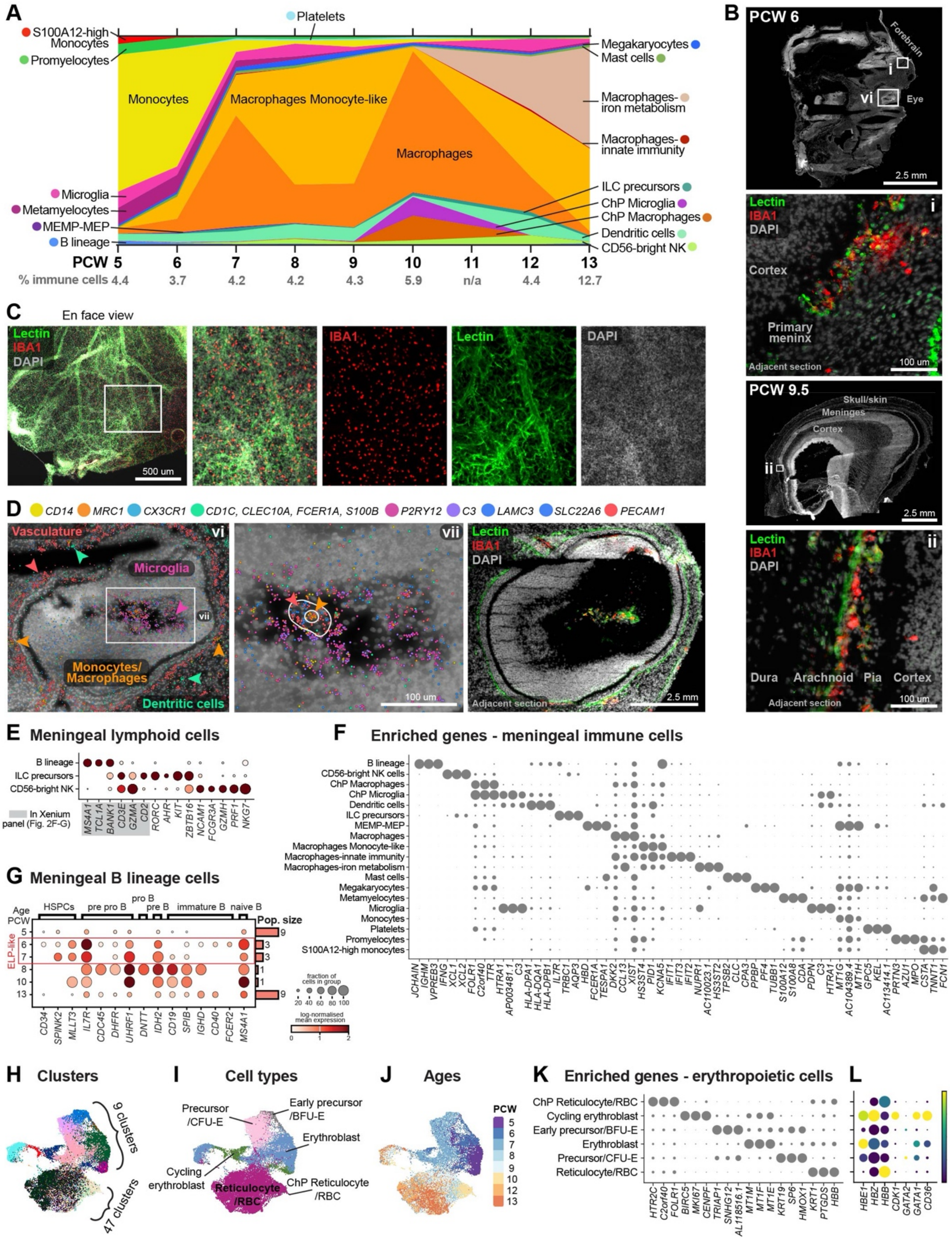
A
S100A12-high Monocytes
Promyelocytes
Platelets
Megakaryocytes
Mast cells
Monocytes
Macrophages Monocyte-like
Macrophages-iron metabolism
Macrophages-innate immunity
Macrophages
Microglia
Metamyelocytes
MEMP-MEP
B lineage
ILC precursors
ChP Microglia
ChP Macrophages
Dendritic cells
CD56-bright NK
PCW 5 6 7 8 9 10 11 12 13
% immune cells 4.4 3.7 4.2 4.2 4.3 5.9 n/a 4.4 12.7
B
PCW 6
Forebrain
Eye
2.5 mm
Lectin
IBA1
DAPI
Cortex
Primary meninx
Adjacent section
100 um
PCW 9.5
Skull/skin
Meninges
Cortex
Dura Arachnoid Pia Cortex
C
En face view
500 um
D
CD14 MRC1 CX3CR1 CD1C, CLEC10A, FCER1A, S100B P2RY12 C3 LAMC3 SLC22A6 PECAM1
Vasculature
Microglia
Monocytes/ Macrophages
Dentritic cells
E Meningeal lymphoid cells
B lineage
ILC precursors
CD56-bright NK
In Xenium panel (Fig. 2F-G)
MS4A1 TCL1A BANK1 CD3E GZMA CD2 RORC AHR KIT ZBTB16 NCAM1 FCGR3A GZMH PRF1 NKG7
F Enriched genes - meningeal immune cells
B lineage
CD56-bright NK cells
ChP Macrophages
ChP Microglia
Dendritic cells
ILC precursors
MEMP-MEP
Macrophages
Macrophages Monocyte-like
Macrophages-innate immunity
Macrophages-iron metabolism
Mast cells
Megakaryocytes
Metamyelocytes
Microglia
Monocytes
Platelets
Promyelocytes
S100A12-high monocytes
G Meningeal B lineage cells
Age PCW
HSPCs pre pro B pro B pre B immature B naive B
ELP-like
Pop. size
CD34 SPINK2 MLLT3 IL7R CDC45 DHFR UHRF1 DNTT IDH2 CD19 SPIB IGHD CD40 FCER2 MS4A1
fraction of cells in group
log-normalised mean expression
H Clusters
9 clusters
47 clusters
I Cell types
Precursor /CFU-E
Early precursor /BFU-E
Erythroblast
Cycling erythroblast
Reticulocyte /RBC
ChP Reticulocyte /RBC
J Ages
PCW
K Enriched genes - erythropoietic cells
ChP Reticulocyte/RBC
Cycling erythroblast
Early precursor/BFU-E
Erythroblast
Precursor/CFU-E
Reticulocyte/RBC
HTR2C C2orf40 FOLR1 BIRC5 MKI67 CENPF TRIAP1 SNHG12 AL118516.1 MT1M MT1F MT1E KRT19 SP6 HMOX1 KRT1 PTGDS HBB
L
HBE1 HBZ HBB CDK1 GATA2 GATA1 CD36

**Figure S2. Macrophages/microglia, lymphoid, and erythropoietic cells, related to Figure 2**
(A) Proportion of immune cell types by age. In grey: proportion of immune cells by age compared to all cells in the meninges.
(B) IBA1 Immunostaining (macrophages and microglia), Lectin dye (vasculature), and DAPI (nuclei) in sagittal sections adjacent to sections used for spatial transcriptomics.
(C) As (B) but PCW 9.5 floating meninges.
(D) PCW 6 eye, with spatial transcriptomic data showing dots representing RNA molecules of markers for monocytes/macrophages, microglia, and dendritic cells, and IBA1 immunostaining.
(E) Dot plot showing the expression of lymphoid markers in a subset of meningeal lymphoid cells. Grey box indicates genes also included in the spatial transcriptomics (Xenium *in situ*) probe panel.
(F) Enriched genes for fetal meningeal immune cells.
(G) Dot plot of scRNAseq gene expression of B-lineage markers in developmental meninges across time (from 5-13 PCW). The size of the dot represents the % of cells within a group that express a given gene, and the colour of the dot indicates ln(x+1) mean gene expression within the group, where x equal counts normalised to 1e4 per cell. ELP, early lymphoid cell.
(H) UMAP of erythropoietic cells coloured by clusters.
(I) UMAP of erythropoietic cells coloured by cell type annotation.
(J) UMAP of erythropoietic cells coloured by sample ages.
(K) Enriched genes per erythropoietic cell annotation.
(L) Dot plot showing expression of yolk sac-derived primitive (*HBE1/HBZ*) and definitive (*HBB*) markers, and classical genes involved in erythropoiesis.

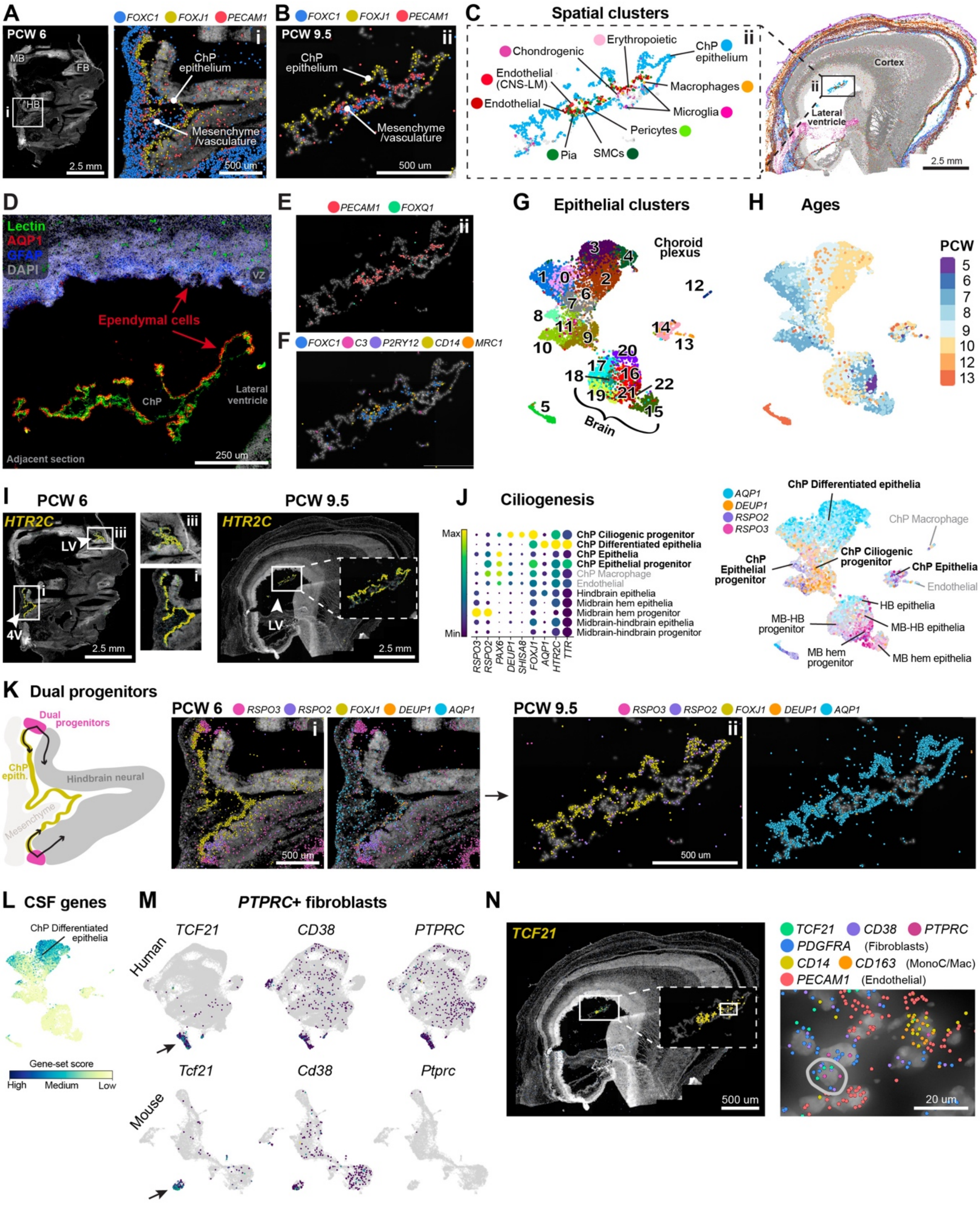
A
PCW 6
FOXC1 FOXJ1 PECAM1
MB
FB
HB
i
2.5 mm
ChP epithelium
Mesenchyme /vasculature
500 um
B
FOXC1 FOXJ1 PECAM1
PCW 9.5
ii
ChP epithelium
Mesenchyme /vasculature
500 um
C
Spatial clusters
Chondrogenic
Erythropoietic
ChP epithelium
Endothelial (CNS-LM)
Endothelial
Macrophages
Microglia
Pericytes
Pia
SMCs
Cortex
Lateral ventricle
2.5 mm
D
Lectin
AQP1
GFAP
DAPI
VZ
Ependymal cells
ChP
Lateral ventricle
Adjacent section
250 um
E
PECAM1 FOXQ1
F
FOXC1 C3 P2RY12 CD14 MRC1
G
Epithelial clusters
Choroid plexus
Brain
H
Ages
PCW
5
6
7
8
9
10
12
13
I
PCW 6
HTR2C
LV
4V
iii
2.5 mm
PCW 9.5
HTR2C
LV
2.5 mm
J
Ciliogenesis
Max
Min
ChP Ciliogenic progenitor
ChP Differentiated epithelia
ChP Epithelia
ChP Epithelial progenitor
ChP Macrophage
Endothelial
Hindbrain epithelia
Midbrain hem epithelia
Midbrain hem progenitor
Midbrain-hindbrain epithelia
Midbrain-hindbrain progenitor
RSPO3
RSPO2
PAX6
DEUP1
SHISA8
FOXJ1
AQP1
HTR2C
TTR
AQP1
DEUP1
RSPO2
RSPO3
ChP Differentiated epithelia
ChP Macrophage
ChP Ciliogenic progenitor
ChP Epithelial progenitor
ChP Epithelia
Endothelial
HB epithelia
MB-HB progenitor
MB-HB epithelia
MB hem progenitor
MB hem epithelia
K
Dual progenitors
Dual progenitors
ChP epith.
Hindbrain neural
Mesenchyme
PCW 6
RSPO3 RSPO2 FOXJ1 DEUP1 AQP1
500 um
PCW 9.5
RSPO3 RSPO2 FOXJ1 DEUP1 AQP1
500 um
L
CSF genes
ChP Differentiated epithelia
Gene-set score
High
Medium
Low
M
PTPRC+ fibroblasts
Human
TCF21
CD38
PTPRC
Mouse
Tcf21
Cd38
Ptprc
N
TCF21
500 um
TCF21 CD38 PTPRC
PDGFRA (Fibroblasts)
CD14 CD163 (MonoC/Mac)
PECAM1 (Endothelial)
20 um

**Figure S3. Fetal choroid plexus development, related to Figure 3**
(A) Spatial transcriptomic data showing *FOXC1*+ mesenchymal/meningeal progenitors, endothelial cells (*PECAM1*), and choroid plexus (ChP) ependymal cells (*FOXJ1*) at PCW 6. FB, forebrain; MB, midbrain; HB, hindbrain.
(B) Same as (A) but at PCW 9.5.
(C) Annotated spatial clusters of the lateral ventricle ChP at PCW 9.5.
(D) Lectin dye (vasculature), DAPI (nuclei) and Aquaporin 1 (AQP1) immunostaining.
(E) Spatial transcriptomic data showing *PECAM1* and *FOXQ1* RNA molecules in the ChP stroma.
(F) Same as (E) but *CD14*/*MRC1*+ monocytes/macrophages, *FOXC1*+ ChP meningeal stroma, and *C3/P2RY12*+ microglia.
(G) scRNA-seq UMAP of ChP and brain epithelial cells, coloured by clusters.
(H) UMAP of epithelial cells coloured by sample ages.
(I) Spatial transcriptomics showing *HTR2C* expression.
(J) Dot plot showing the expression of genes involved in epithelial ciliogenesis (left), per annotated cell type on UMAP (right).
(K) Schematic and spatial validation of *RSPO2/3*+ dual neuro-epithelial progenitors, and ciliogenesis markers, at PCW 6 and 9.5.
(L) UMAP of epithelial cells, coloured by a CSF gene-set score (Methods).
(M) UMAPs of human fetal (this study) and mouse embryonic[41] meningeal fibroblasts coloured by expression of *TCF21/Tcf21*, *PTPRC/Ptprc*, and *CD38/Cd38* on a grey background of all cells.
(N) Spatial validation of *TCF21*+ *PTPRC*+ *CD38*+ fibroblasts in the ChP.

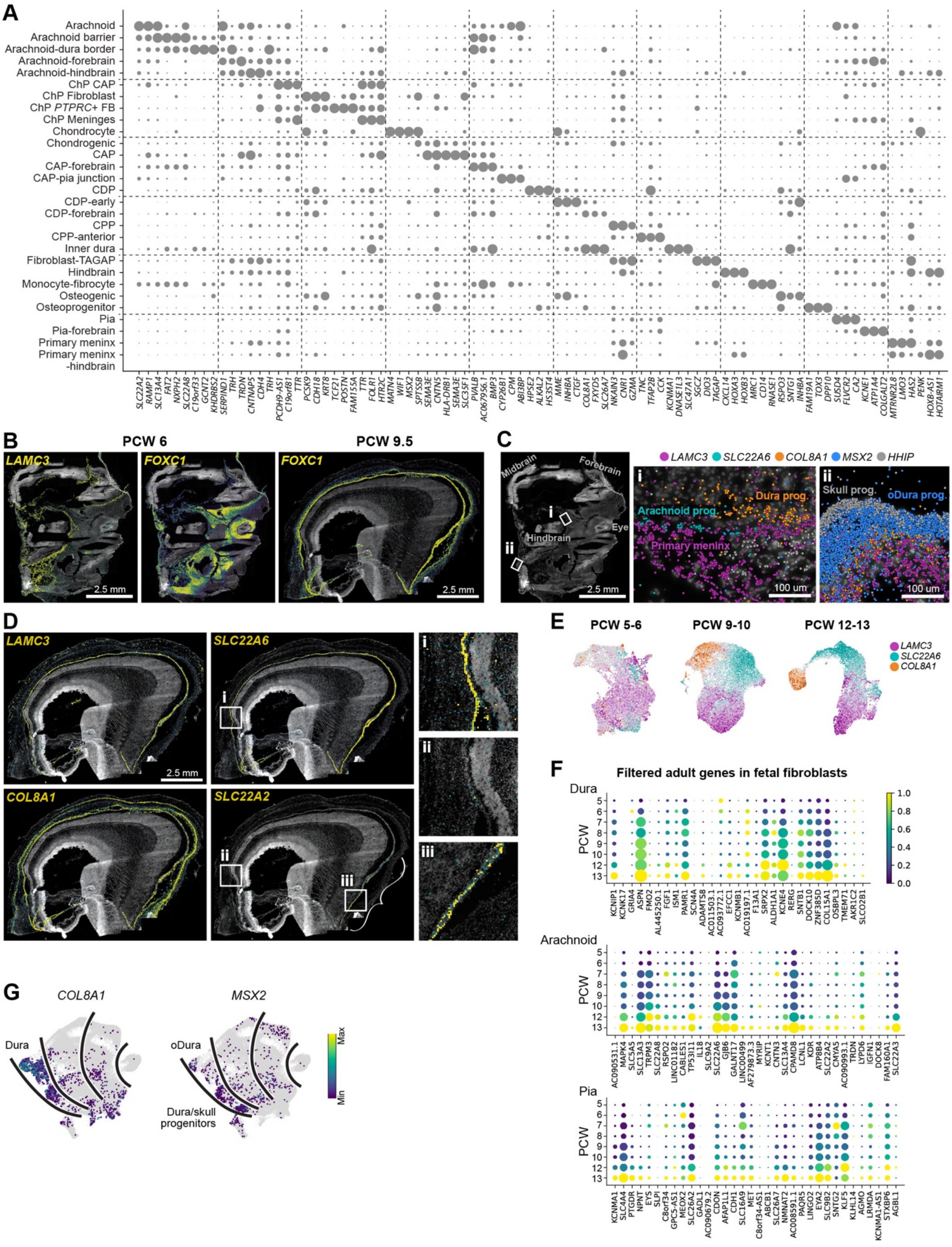
A
Arachnoid
Arachnoid barrier
Arachnoid-dura border
Arachnoid-forebrain
Arachnoid-hindbrain
ChP CAP
ChP Fibroblast
ChP PTPRC+ FB
ChP Meninges
Chondrocyte
Chondrogenic
CAP
CAP-forebrain
CAP-pia junction
CDP
CDP-early
CDP-forebrain
CPP
CPP-anterior
Inner dura
Fibroblast-TAGAP
Hindbrain
Monocyte-fibrocyte
Osteogenic
Osteoprogenitor
Pia
Pia-forebrain
Primary meninx
Primary meninx -hindbrain
B
PCW 6
PCW 9.5
LAMC3
FOXC1
2.5 mm
C
Midbrain
Forebrain
Hindbrain
Eye
LAMC3 SLC22A6 COL8A1 MSX2 HHIP
Dura prog.
Arachnoid prog.
Primary meninx
Skull prog.
oDura prog.
100 um
D
LAMC3
SLC22A6
COL8A1
SLC22A2
E
PCW 5-6
PCW 9-10
PCW 12-13
F
Filtered adult genes in fetal fibroblasts
Dura
Arachnoid
Pia
PCW
G
COL8A1
MSX2
Dura
oDura
Dura/skull progenitors
Max
Min

**Figure S4. Meningeal layer marker dynamics, related to Figure 4**
(A) Dotplot of enriched genes per CellType annotation. CAP, Committed arachnoid precursor; CDP, Committed dura precursor; ChP, Choroid plexus; CPP, Committed pia precursor.
(B) Spatial transcriptomics showing *LAMC3* and *FOXC1* expression at PCW 6, and *FOXC1* at 9.5.
(C) Spatial transcriptomics of identified layer markers *LAMC3* (pia), *SLC22A6* (arachnoid), *COL8A1* (dura), *MSX2* (outer dura, skull and periosteum), and *HHIP* (skull) at PCW 6.
(D) Spatial transcriptomics showing *LAMC3, SLC22A6, COL8A1,* and *SLC22A2* expression at PCW 9.5.
(E) UMAPs of subsets of pooled PCW 5-6, 9-10, and 12-13 fibroblasts. Cells are coloured by their expression of the gene they express most highly among *LAMC3* (pia), *SLC22A6* (arachnoid), *COL8A1* (dura).
(F) Dotplot showing the expression of enriched genes from adult pia, arachnoid, and dura (Fig. 4F), in fetal fibroblasts over developmental timepoints.
(G) UMAP showing the expression of *COL8A1* (dura) and *MSX2* (outer dura) in fetal meningeal fibroblasts.

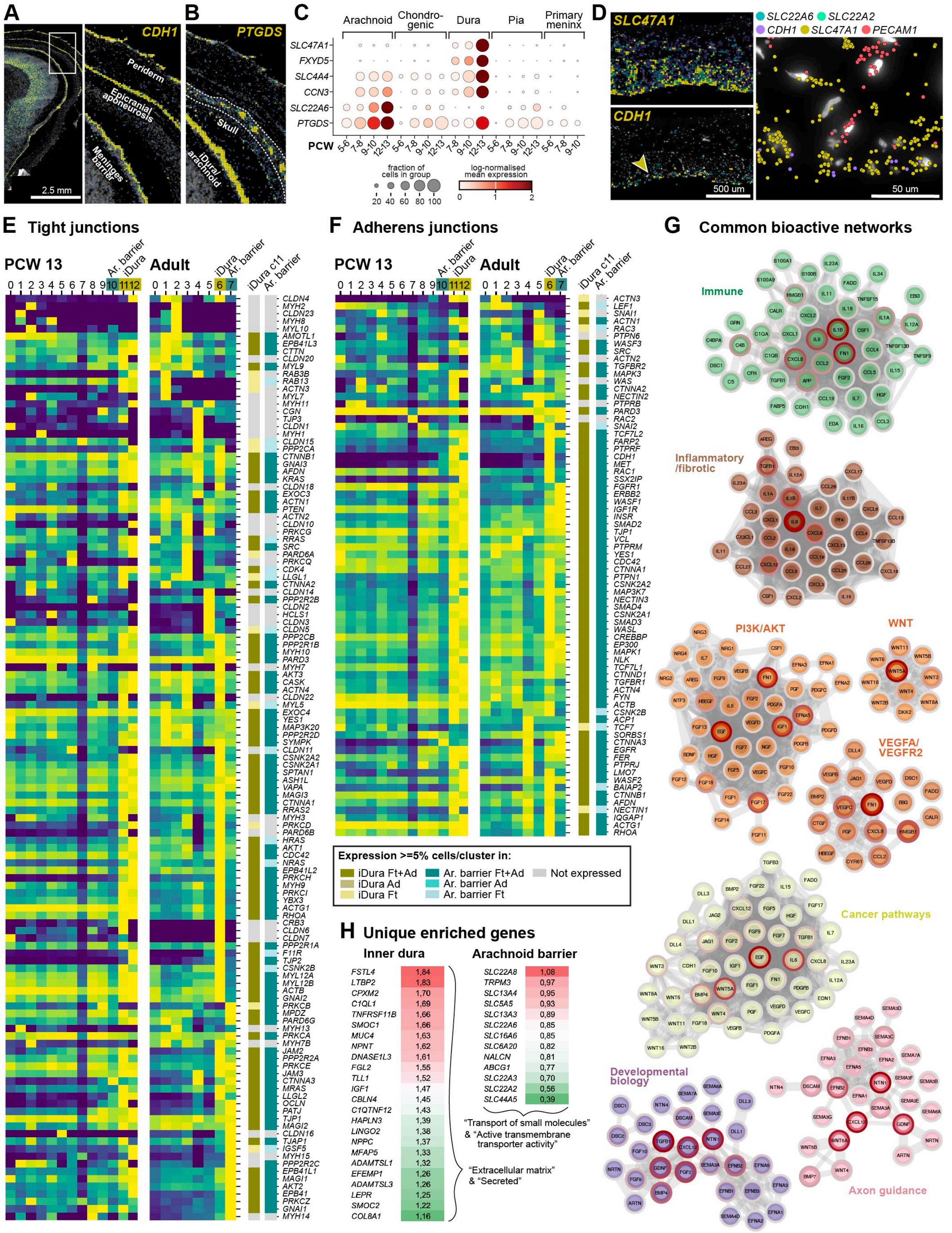

A
CDH1
Periderm
Epicranial aponeurosis
Meninges barrier
2.5 mm
B
PTGDS
Skull
iDura/ arachnoid
C
Arachnoid
Chondro-genic
Dura
Pia
Primary meninx
SLC47A1
FXYD5
SLC4A4
CCN3
SLC22A6
PTGDS
PCW
fraction of cells in group
20 40 60 80 100
log-normalised mean expression
0 1 2
D
SLC47A1
CDH1
SLC22A6
SLC22A2
SLC47A1
PECAM1
500 um
50 um
E Tight junctions
PCW 13
Adult
Ar. barrier
iDura
iDura c11
F Adherens junctions
Expression >=5% cells/cluster in:
iDura Ft+Ad
iDura Ad
iDura Ft
Ar. barrier Ft+Ad
Ar. barrier Ad
Ar. barrier Ft
Not expressed
G Common bioactive networks
Immune
Inflammatory /fibrotic
PI3K/AKT
WNT
VEGFA/ VEGFR2
Cancer pathways
Developmental biology
Axon guidance
H Unique enriched genes
Inner dura
FSTL4 1,84
LTBP2 1,83
CPXM2 1,70
C1QL1 1,69
TNFRSF11B 1,66
SMOC1 1,66
MUC4 1,63
NPNT 1,62
DNASE1L3 1,61
FGL2 1,55
TLL1 1,52
IGF1 1,47
CBLN4 1,45
C1QTNF12 1,43
HAPLN3 1,39
LINGO2 1,38
NPPC 1,37
MFAP5 1,33
ADAMTSL1 1,32
EFEMP1 1,26
ADAMTSL3 1,26
LEPR 1,25
SMOC2 1,22
COL8A1 1,16
Arachnoid barrier
SLC22A8 1,08
TRPM3 0,97
SLC13A4 0,95
SLC5A5 0,93
SLC13A3 0,89
SLC22A6 0,85
SLC16A6 0,85
SLC6A20 0,82
NALCN 0,81
ABCG1 0,77
SLC22A3 0,70
SLC22A2 0,56
SLC44A5 0,39
"Transport of small molecules" & "Active transmembrane transporter activity"
"Extracellular matrix" & "Secreted"

**Figure S5. Shared and unique gene expression in inner dura and arachnoid barrier cells, related to Figure 5**

(A) Spatial transcriptomics showing *CDH1* expression in PCW 9.5 head fibroblasts.
(B) Same as (A) but *PTGDS* expression.
(C) Dotplot showing the expression of arachnoid and dura-specific genes over time, in arachnoid, chondrogenic, dura, pia, and primary meninx.
(D) Spatial transcriptomics showing *SLC47A1*, *CDH1*, *SLC22A2*, *SLC22A6* and *PECAM1* expression in human adult dura.
(E) Heatmap showing the expression of genes from the full KEGG list of tight junction genes (#M11355), in scRNA-seq clusters of PCW 13 and adult meningeal fibroblast. Right: Stylised heatmaps with a threshold criteria of genes being expressed in at least 5% of cells/cluster.
(F) Same as (E) but the full KEGG list of adherens junction genes (#M638).
(G) STRING interaction networks of shared bioactive genes between inner dura and arachnoid barrier clusters, split into subsets of significantly enriched functional pathways. Gene centrality (connectedness) outlined by red.
(H) Subset of enriched genes in inner dura and arachnoid barrier clusters, and relevant ontologies.

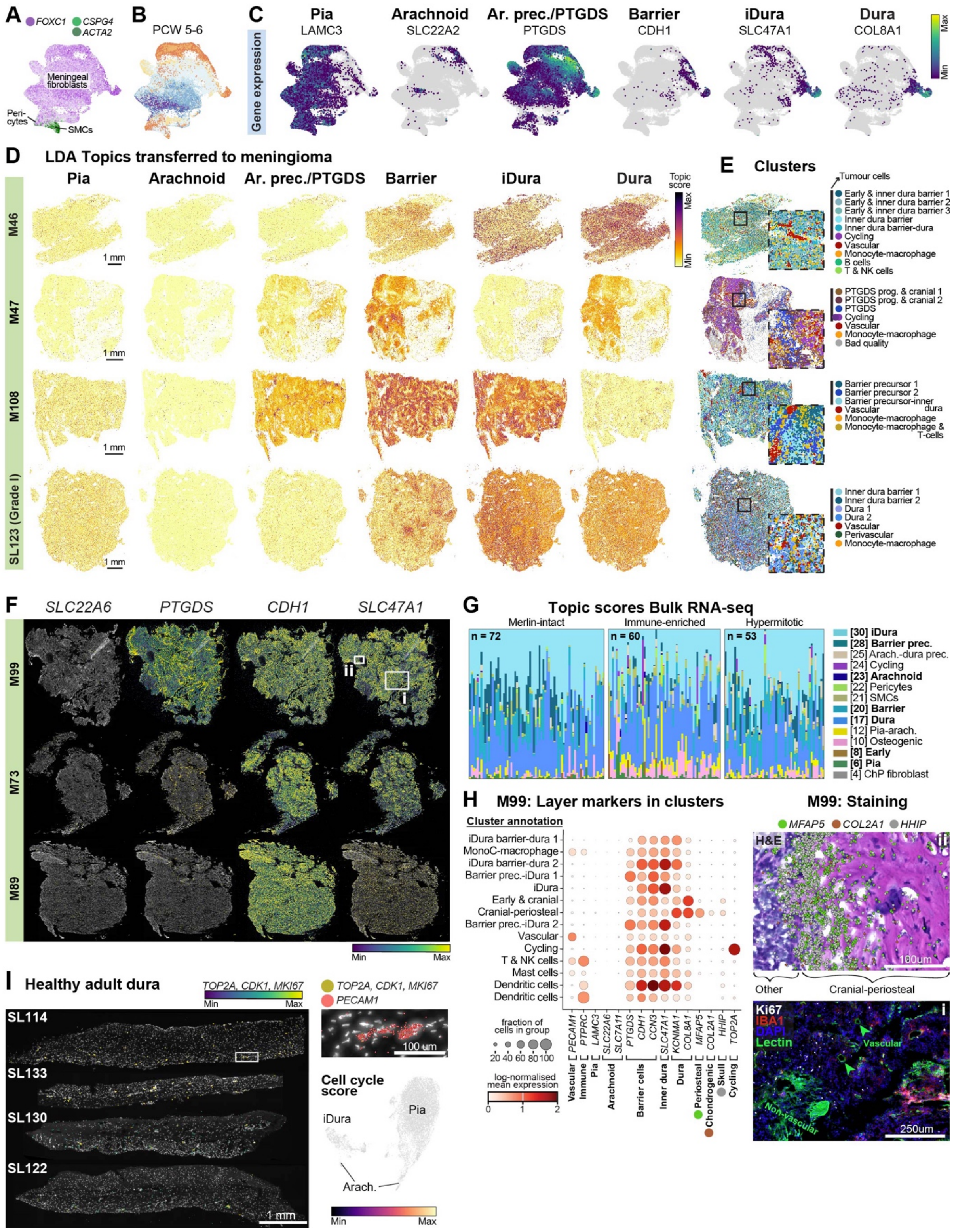

A
FOXC1
CSPG4
ACTA2
Meningeal fibroblasts
Peri-cytes
SMCs
B
PCW 5-6
C
Gene expression
Pia
LAMC3
Arachnoid
SLC22A2
Ar. prec./PTGDS
PTGDS
Barrier
CDH1
iDura
SLC47A1
Dura
COL8A1
Max
Min
D
LDA Topics transferred to meningioma
Pia
Arachnoid
Ar. prec./PTGDS
Barrier
iDura
Dura
Topic score
M46
M47
M108
SL123 (Grade I)
1 mm
E
Clusters
Tumour cells
Early & inner dura barrier 1
Early & inner dura barrier 2
Early & inner dura barrier 3
Inner dura barrier
Inner dura barrier-dura
Cycling
Vascular
Monocyte-macrophage
B cells
T & NK cells
PTGDS prog. & cranial 1
PTGDS prog. & cranial 2
PTGDS
Cycling
Vascular
Monocyte-macrophage
Bad quality
Barrier precursor 1
Barrier precursor 2
Barrier precursor-inner dura
Vascular
Monocyte-macrophage
Monocyte-macrophage & T-cells
Inner dura barrier 1
Inner dura barrier 2
Dura 1
Dura 2
Vascular
Perivascular
Monocyte-macrophage
F
SLC22A6
PTGDS
CDH1
SLC47A1
M99
M73
M89
ii
i
Min
Max
G
Topic scores Bulk RNA-seq
Merlin-intact
Immune-enriched
Hypermitotic
n = 72
n = 60
n = 53
[30] iDura
[28] Barrier prec.
[25] Arach.-dura prec.
[24] Cycling
[23] Arachnoid
[22] Pericytes
[21] SMCs
[20] Barrier
[17] Dura
[12] Pia-arach.
[10] Osteogenic
[8] Early
[6] Pia
[4] ChP fibroblast
H
M99: Layer markers in clusters
Cluster annotation
iDura barrier-dura 1
MonoC-macrophage
iDura barrier-dura 2
Barrier prec.-iDura 1
iDura
Early & cranial
Cranial-periosteal
Barrier prec.-iDura 2
Vascular
Cycling
T & NK cells
Mast cells
Dendritic cells
Dendritic cells
PECAM1
PTPRC
LAMC3
SLC22A6
SLC7A11
PTGDS
CDH1
CCN3
SLC47A1
KCNMA1
COL8A1
MFAP5
COL2A1
HHIP
TOP2A
Vascular
Immune
Pia
Arachnoid
Barrier cells
Inner dura
Dura
Periosteal
Chondrogenic
Skull
Cycling
fraction of cells in group
20 40 60 80 100
log-normalised mean expression
0 1 2
M99: Staining
MFAP5
COL2A1
HHIP
H&E
ii
100um
Other
Cranial-periosteal
Ki67
IBA1
DAPI
Lectin
i
Vascular
Non-vascular
250um
I
Healthy adult dura
TOP2A, CDK1, MKI67
Min
Max
SL114
SL133
SL130
SL122
1 mm
TOP2A, CDK1, MKI67
PECAM1
100 um
Cell cycle score
iDura
Pia
Arach.
Min
Max

**Figure S6. Further analysis demonstrating that meningioma tumours are dura-like, related to Figure 6**

(A) Cells coloured by their expression of the gene they express most highly among *FOXC1* (meningeal fibroblasts & pericytes), *CSPG4* (pericytes), *ACTA2* (SMCs).
(B) UMAP of fetal fibroblasts and perivascular cells coloured by sample ages.
(C) Genes ranked highly by our LDA model for meningeal layer topics. Gene expression shown on UMAP of fetal fibroblasts and perivascular cells, on grey background of all cells. Ar. prec., Arachnoid precursors; iDura, inner dura.
(D) LDA topics transferred to spatial data from three high grade hypermitotic meningioma tumours; M46, M47 & M108, and one grade I tumour; SL123.
(E) Spatial clusters of tumours shown in (D), coloured by their most similar meningeal layer.
(F) Spatial transcriptomic data showing *SLC22A6*, *PTGDS*, *CDH1*, and *SLC47A1* expression in M99, M73, and M89 tumours. Inset references i and ii are for (H).
(G) Stacked bar chart showing scores of LDA topics transferred to bulk RNA-sequencing data from 185 meningioma tumours[18].
(H) Left: Dot plot showing gene expression of meningeal layer markers in M99 clusters. MonoC, Monocytes. Top right: Spatial transcriptomic data of *MFAP5*, *COL2A1* and *HHIP*, on H&E staining. Bottom right: Immunohistochemistry of Ki67, and IBA1, and DAPI and Lectin dyes. Inset references i and ii can be seen in (F).
(I) Spatial transcriptomic data of four human adult dura samples, showing RNA molecules of cycling cells (*TOP1A, CDK1, MKI67*) and endothelial cells (*PECAM1*). Cell cycle score on scRNA-seq UMAP of adult fibroblasts.

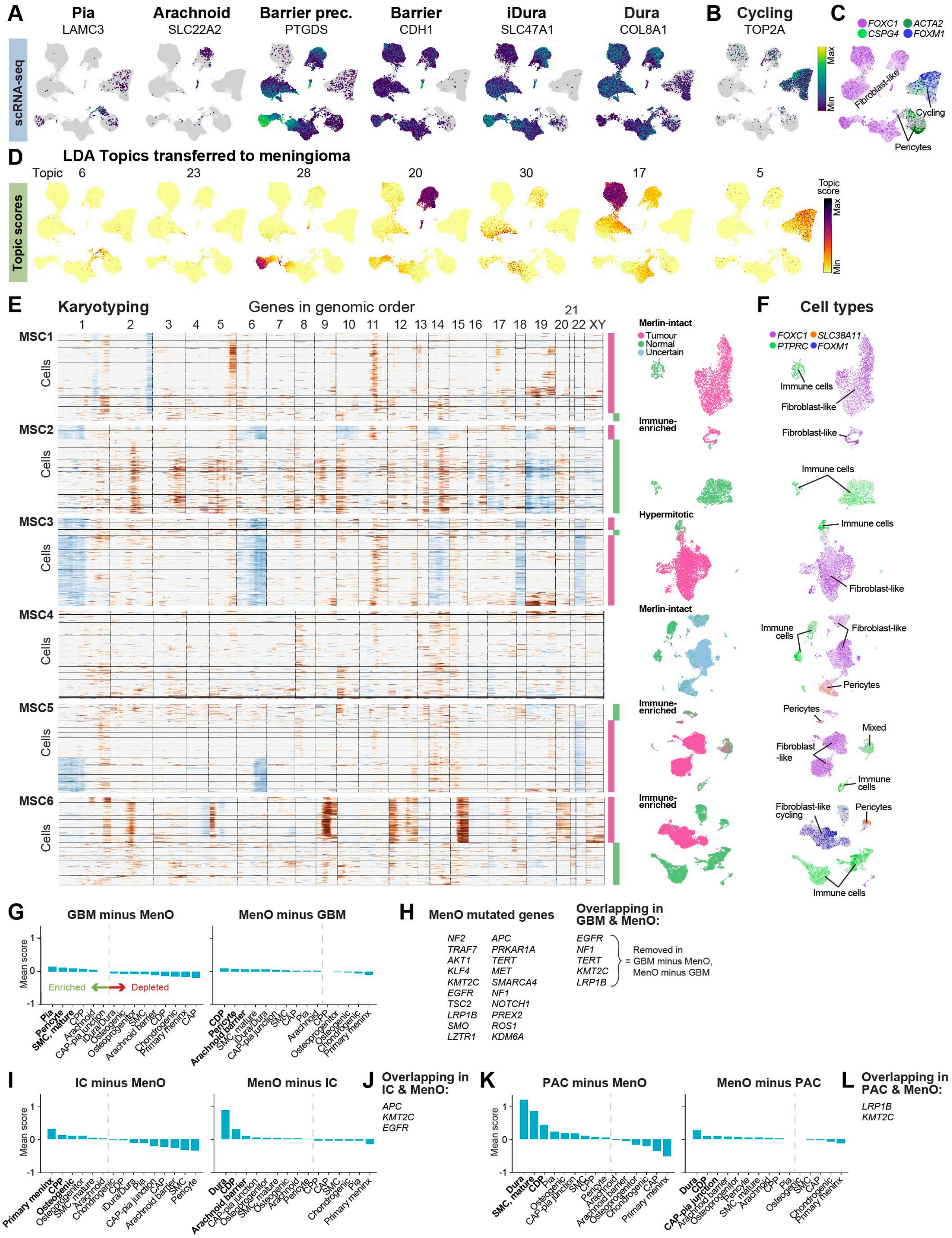
A
Pia LAMC3
Arachnoid SLC22A2
Barrier prec. PTGDS
Barrier CDH1
iDura SLC47A1
Dura COL8A1
scRNA-seq
B Cycling TOP2A
Max
Min
C FOXC1 ACTA2 CSPG4 FOXM1
Fibroblast-like
Cycling
Pericytes
D LDA Topics transferred to meningioma
Topic 6 23 28 20 30 17 5
Topic scores
Topic score
E Karyotyping
Genes in genomic order
MSC1
MSC2
MSC3
MSC4
MSC5
MSC6
Cells
Merlin-intact
Tumour
Normal
Uncertain
Immune-enriched
Hypermitotic
F Cell types
FOXC1 SLC38A11 PTPRC FOXM1
Immune cells
Fibroblast-like
Pericytes
Mixed
Fibroblast-like cycling
G GBM minus MenO
MenO minus GBM
Mean score
Enriched
Depleted
H MenO mutated genes
NF2 APC
TRAF7 PRKAR1A
AKT1 TERT
KLF4 MET
KMT2C SMARCA4
EGFR NF1
TSC2 NOTCH1
LRP1B PREX2
SMO ROS1
LZTR1 KDM6A
Overlapping in GBM & MenO:
EGFR
NF1
TERT
KMT2C
LRP1B
= Removed in GBM minus MenO, MenO minus GBM
I IC minus MenO
MenO minus IC
J Overlapping in IC & MenO:
APC
KMT2C
EGFR
K PAC minus MenO
MenO minus PAC
L Overlapping in PAC & MenO:
LRP1B
KMT2C

**Figure S7. Analysis of meningioma scRNA-seq, and mutated genes from various cancers in fetal meningeal fibroblasts, related to Figure 7**

(A) UMAP of tumour cells, coloured by the expression of genes ranked highly by our LDA model for meningeal layer topics, on a grey background of all cells.
(B) UMAP of tumour cells, coloured by *TOP2A* expression.
(C) Cells coloured by their expression of the gene they express most highly among *FOXC1* (meningeal fibroblasts & pericytes), *CSPG4* (pericytes), *ACTA2* (SMCs), and *FOXM1* (cycling tumour).
(D) LDA topics for meningeal layers and cycling cells (Topic 5) transferred to meningioma scRNA-seq data.
(E) Karyotyping of individual tumours. Chromosomal deletions are blue, and gains are brown. Bars to the right and UMAPs of individual tumours indicate cells that are tumour (pink), normal (green) and of uncertain karyotype (light blue).
(F) Annotation of individual tumour samples, and expression of *FOXC1* (meningeal), *SLC38A11* (pericytes), *ACTA2* (SMCs), *PTPRC* (immune).
(G) Quantification of gene-set score of top 20 mutated genes in glioblastoma (GBM) tumours minus genes overlapping with top 20 mutated genes in meningioma (MenO), and MenO minus GBM overlapping genes, in fetal meningeal fibroblast and perivascular cell Subclass (as in Figure 7F).
(H) Top 20 mutated genes in meningioma, and genes overlapping between the top 20 mutated genes in GBM and MenO.
(I) Same as (G) but for intestinal carcinomas (IC).
(J) List of overlapping genes between the top 20 mutated genes in IC and MenO.
(K) Same as (G) but for prostate adenocarcinomas (PAC).
(L) List of overlapping genes between the top 20 mutated genes in PAC and MenO.

**Table S1.** Samples

**Table S2.** scRNA-seq cluster metadata

**Table S3.** Xenium probe panels

**Table S4.** Xenium clusters

**Table S5.** CSF analysis, related to Figure S3

**Table S6.** Network analysis, related to Figures 5 and S5

**Table S7.** Topics filtered genes, related to Figures 6, S6, and S7

**Table S8.** COSMIC mutated genes, related to Figure 7 and S7
